# *Austropuccinia psidii*, causing myrtle rust, has a gigabase-sized genome shaped by transposable elements

**DOI:** 10.1101/2020.03.18.996108

**Authors:** Peri A Tobias, Benjamin Schwessinger, Cecilia H Deng, Chen Wu, Chongmei Dong, Jana Sperschneider, Ashley Jones, Zhenyan Lou, Peng Zhang, Karanjeet Sandhu, Grant R Smith, Josquin Tibbits, David Chagné, Robert F Park

**Affiliations:** School of Life and Environmental Sciences, University of Sydney, Camperdown, NSW 2006, Australia; Plant & Food Research Australia, Sydney, Australia; Australia Research School of Biology, The Australian National University, Acton, ACT 2601; The New Zealand Institute for Plant and Food Research Limited, Private Bag 92169, Auckland 1142, New Zealand; Plant Breeding Institute, University of Sydney, Private Bag 4011 Narellan, NSW 2567, Australia; Biological Data Science Institute, The Australian National University, Canberra, ACT, 2600, Australia; The New Zealand Institute for Plant and Food Research Limited, Private Bag 4704, Christchurch Mail Centre, Christchurch 8140, New Zealand; Agriculture Victoria Department of Jobs, Precincts and Regions, Bundoora, Vic 3083, Australia; The New Zealand Institute for Plant & Food Research, Private Bag 11600, Palmerston North 4442, New Zealand

**Keywords:** myrtle rust, Pucciniomycotina, fungal genome evolution, Myrtaceae, transposable elements

## Abstract

*Austropuccinia psidii*, originating in South America, is a globally invasive fungal plant pathogen that causes rust disease on Myrtaceae. Several biotypes are recognized, with the most widely distributed pandemic biotype spreading throughout the Asia-Pacific and Oceania regions over the last decade. *Austropuccinia psidii* has a broad host range with more than 480 myrtaceous species. Since first detected in Australia in 2010, the pathogen has caused the near extinction of at least three species and negatively affected commercial production of several Myrtaceae. To enable molecular and evolutionary studies into *A. psidii* pathogenicity, we assembled a highly contiguous genome for the pandemic biotype. With an estimated haploid genome size of just over 1 Gb (gigabases), it is the largest assembled fungal genome to date. The genome has undergone massive expansion via distinct transposable element (TE) bursts. Over 90% of the genome is covered by TEs predominantly belonging to the Gypsy superfamily. These TE bursts have likely been followed by deamination events of methylated cytosines to silence the repetitive elements. This in turn led to the depletion of CpG sites in transposable elements and a very low overall GC content of 33.8%. The overall gene content is highly conserved, when compared to other closely related Pucciniales, yet the intergenic distances are increased by an order of magnitude indicating a general insertion of TEs between genes. Overall, we show how transposable elements shaped the genome evolution of *A. psidii* and provide a greatly needed resource for strategic approaches to combat disease spread.

## Introduction

The globally invasive fungal plant pathogen, *Austropuccinia psidii* (G. Winter) Beenken (Beenken 2017), was first reported in South America (Winter 1884) on guava (*Psidium guajava*) and named *Puccinia psidii*. This pathogen is the causal agent of rust disease on Myrtaceae, with guava rust, eucalyptus rust, ‘ōhi’a rust and myrtle rust used as disease names. Several biotypes are recognized (Stewart et al. 2018; Kim et al. 2017) with only the pandemic biotype (Machado et al. 2015) currently believed to be present in the Asia-Pacific and Oceania regions (du Plessis et al. 2019; Sandhu et al. 2015). The disease represents a relatively recent arrival in these geographic regions with first detection in Hawaii in 2005, China in 2009, Australia in 2010, New Caledonia in 2013, and New Zealand in 2017 (Carnegie & Pegg 2018). It appears that *A. psidii* is spreading rapidly and causing devastating impacts to natural vegetation communities (Soewarto et al. 2018; Carnegie et al. 2015). *Austropuccinia psidii* is a biotrophic pathogen and has a macrocyclic life cycle but predominates in the asexual, dikaryotic urediniospore state with two haploid nuclei in each cell (Coutinho et al. 1998). Germination of wind-borne urediniospores requires the presence of free water, darkness and optimal temperatures between 15 and 25 °C, with cuticular penetration occurring in susceptible hosts within 12 hours (Hunt 1968). During extended cold periods (∼8 °C) the pathogen can overwinter in a dormant state within the host plant (Beresford et al. 2020). The pathogen causes disease symptoms on the new foliage, stems and buds (Tobias et al. 2015; Carnegie et al. 2015) of a wide range of perennial plants with over 480 known host species globally (Soewarto et al. 2019). Repeated infections can lead to crown loss and plant mortality (Pegg et al. 2017). Globally, there are 5,950 described myrtaceous species (Christenhusz & Byng 2016), of which 2,250 are endemic to Australia, where myrtle rust has been particularly damaging (Berthon et al. 2018). For example, two previously wide-spread east Australian plant species, *Rhodamnia rubescens* (Benth.) Miq. and *Rhodomyrtus psidioides* (G.Don) Benth., and one recently described species, *Lenwebbia* sp. Main Range (P.R.Sharpe 4877), have now been listed as critically endangered owing to repeated infection causing death (Pegg et al. 2017; Makinson 2018). Myrtle rust has also caused the decline of at least one keystone species, *Melaleuca quinquenervia* (Cav.) S.T.Blake (Pegg et al. 2017), and affected commercial production of Myrtaceae such as tea tree (*Melaleuca alternifolia* (Maiden & Betche) Cheel) and lemon myrtle (*Backhousia citriodora* F. Muell.) (Carnegie & Pegg 2018). It is currently unclear what drives the broad host range of *A. psidii* which is atypical of rust fungi. There is little understanding of how this pathogen overcomes defense mechanisms and infects its diverse range of host plants, and even less understanding of why widespread, but very variable, within host resistance is observed.

We generated a high-quality reference genome of the pandemic biotype of *A. psidii* using a combination of long-read sequencing and Hi-C technology (van Berkum et al. 2010). Our final haploid genome assembly for *A. psidii* was 1.02 Gb (of an estimated 1.0 to 1.4 Gb based on flow cytometry and k-mer analysis, this study) with 66 contigs and 29 telomeres. This is a significant improvement over previous draft assemblies based on short reads (Tan et al. 2014) or linked-read sequence data (McTaggart et al. 2018), which were either unreasonably small (<150 Mbp) or fragmented (more than 100,000 contigs) highlighting the difficulty of assembling this genome. In general, dikaryotic genomes of rust fungi are known to be rich in transposable elements (TE) and have high levels of heterozygosity (Schwessinger et al. 2018; Zheng et al. 2013). The *A. psidii* genome further illustrates this, being the largest assembled fungal genome to date. Clearly, transposable elements have dramatically shaped evolution of the *A*. *psidii* genome, which might have contributed to the expansion of its host range. The high contiguity of the genome, combined with the unique genomic features identified in this study, provide essential resources to understand and address the spread and evolution of this invasive plant pathogen.

## Results

### Flow cytometry and k-mer analysis predict a haploid genome size of 1Gbp

To guide the quantity of long read sequence required to assemble the *A. psidii* genome, the nuclear DNA content of *A. psidii* cells was estimated using flow cytometry. We estimated the 1C (single nucleus) genome content at 1.42 pg (1,388.91 Mbp) using a method for biotrophic fungi (Tavares et al. 2014) and the host genome (*Eucalyptus grandis*) as the internal reference. As an additional approach, we ran a k-mer analysis and predicted the diploid genome size at 2,100 Mbp. As the initial data included mitochondrial sequences, we proceeded with an estimated haploid genome of 1 Gbp. We initially sequenced and assembled reads from 21 PacBio SMRT cells and, after deeming the coverage level inadequate (Table 1), sequenced a further eight cells (18X RSII and 11X Sequel).

### Long-read genome assembly leads to a 2 Gbp partially phased di-haploid genome

We obtained 162 Gigabytes (GB) of data from 29 SMRT cells. An initial diploid assembly with Canu (v1.6) (Koren et al. 2016) using data from 21 SMRT cells (36.1 X coverage) produced an assembly size of 1.52 Gbp (Table 1 A). The contig numbers for this assembly were high and completeness was low at 79.8% complete and 7.0% fragmented, based on Benchmarking Universal Single-Copy Orthologs (BUSCO) in genome mode (Simão et al. 2015). We therefore further increased the genome coverage for PacBio long reads (72.4 X coverage) and assembled the di-haploid genome of *A. psidii*. The final assembly size was 1.98 Gb, with 13,361 contigs (Table 1 A).

**Table 1.** (A) Comparative statistics for the *Austropuccinia psidii* raw sequence and assembly at different sequencing depth. (B) Partially phased primary and secondary assemblies after deduplication with Purge Haplotigs (Roach et al. 2018) before, (C) after Hi-C scaffolding.

| (A) Raw data statistics |  | 21 SMRT cells |  |  | 29 SMRT cells |  |
| --- | --- | --- | --- | --- | --- | --- |
| Raw read numbers |  | 2 696 004 |  |  | 5 798 431 |  |
| Raw bases |  | 36 099 416 447 |  |  | 72 410 490 565 |  |
| Coverage (X) |  | 36.1 |  |  | 72.4 |  |
| Corrected mean length (bp) |  | 11 733 |  |  | 21 811 |  |
| Corrected N50 (bp) |  | 47 581 |  |  | 46 751 |  |
| Assembly statistics |  |  |  |  |  |  |
| Assembly parameters |  | diploid |  |  | diploid |  |
| Number of contigs |  | 22 474 |  |  | 13 361 |  |
| Total length (bp) |  | 1 520 827 096 |  |  | 1 985 401 333 |  |
| N50 (bp) |  | 96 784 |  |  | 287 498 |  |
| Shortest contig |  | 1 004 |  |  | 1 003 |  |
| Longest contig |  | 1 037 919 |  |  | 2 557 042 |  |
| Assembly time (aprox.) |  | 2.5 months |  |  | 4.5 months |  |
| (B) Assembly | Contigs | N50 | L50 | Largest | Bases | GC% |
| Primary | 3 187 | 520 311 | 619 | 2 559 506 | 1 018 086 732 | 33.80 |
| Secondary | 8 626 | 159 727 | 1794 | 957 974 | 933 887 333 | 34.26 |
| (C) | Scaffolds | N50 | L50 | Largest | Bases | GC% |
| Primary | 66 | 56 243 252 | 7 | 124 273 364 | 1 018 398 822 | 33.80 |
| Secondary | 67 | 52 409 407 | 7 | 89 073 602 | 934 744 333 | 34.26 |

We deduplicated the assembly contigs with the Purge Haplotigs pipeline (Roach et al. 2018) to generate a partially phased assembly with primary contigs and secondary haplotigs of ∼ 1 Gbp each. We then scaffolded the primary contigs and haplotigs separately using Hi-C data (Table 1 B and C). With no current cytological evidence for the number of *A. psidii* haploid chromosomes to guide scaffolding, we made predictions for scaffold grouping based on telomeres and evidence from a related rust fungus, *Puccinia graminis* f. sp. *tritici* (*n* = 18) (Boehm et al. 1992). Initially, we produced 40 primary scaffolds but found several telomeres from the pre-scaffolded assembly embedded within scaffolds. Previous studies indicate that telomere localisation in the nucleus show affinity to the nuclear membrane (Laberthonnière et al. 2019), therefore cross-linking during our Hi-C library preparation may have proximity-ligated adjacent telomeres leading to errors in scaffolding. Manual curation before and after scaffolding corrected for this problem by identifying and breaking at the start/end of telomeres for a final number of 66 primary (67 secondary) scaffolds.

We estimated completeness of the gene space with BUSCO (Simão et al. 2015) and identified 94.7% conserved proteins for the primary assembly and 97.6% for the combined primary and secondary assemblies when applied to the final predicted proteome. Overall, we generated a high quality near-complete and partially haplotype phased genome for *A. psidii* that is in agreement with our empirically estimated genome size and displays excellent completeness of the predicted proteome.

### The *A. psidii* genome size expansion is driven by transposable elements

Analysis of the *A. psidii* genome assembly provides evidence for very large and diverse repetitive regions that constitute greater than 91% (932,575,815 bp) of the genome. Initial repeat-masking of the *A. psidii* primary and secondary assemblies with published fungal and viridiplantae repeat databases in RepBase (Bao et al. 2015), showed very low percentages of matches at 19.63% (19.89%) and 6.97% (6.99%) respectively. The A. *psidii* genome is largely made of novel repeat regions prompting further analysis to characterise transposable elements (TEs).

Our TE characterization, with comprehensive *de novo* repeat identification (Flutre et al. 2011), revealed massive repeat expansion in *A. psidii*, largely caused by Class I retrotransposons belonging to the Gypsy superfamily (Figure 1). Most of the expansion was driven by a limited number of TE families expanding at discrete time points (Figure 1 C). For example, the extensive expansion observed around 78% family level identity was mostly caused by a single TE family. This is similar to early expansions (30-50% family level identity), which were driven by only a limited number of TE families. In contrast to this pattern the most recent expansion (>75% family level identity), appears to be driven by a more diverse set of TE families including several Class II superfamilies, such as those belonging to the terminal inverted repeats (TIR) order.

*Most of the prominent TEs from the early expansion (30-50% family level identity) were previously annotated Gypsy elements from P. striiformis* f. sp. *tritici*, *P. graminis* f. sp. *tritici* and *Melampsora larici*☐*populina* found in RepBase (Bao et al. 2015). In contrast, the most recent TE expansion (>75% family level identity) was made up of *de novo* identified TEs using the REPET pipeline (Additional Table 1A). We used the TE reference sequences of the 28 most abundant TE families from *A. psidii* to search related published rust genomes (Figure 6) using RepeatMasker (Smit et al. 1996). In general, we identified the longest TE fragments, the most TE copies, and greatest TE coverage in species from which the TEs were originally identified (Additional Table 1A). Using RepeatMasker, we found very little cross-annotation of TEs beyond the genus level. For example, TEs from the three wheat rust fungi were only found in other Puccinia species but not in *Melampsora larici*☐*populina.* Similarly, *de novo* identified TEs from *A. psidii* were not identified in any other rust species at high copy numbers, or with long copies, indicating that they were not high in abundance in the most recent common ancestor and have expanded specifically in *A. psidii*.

**Fig. 1.**
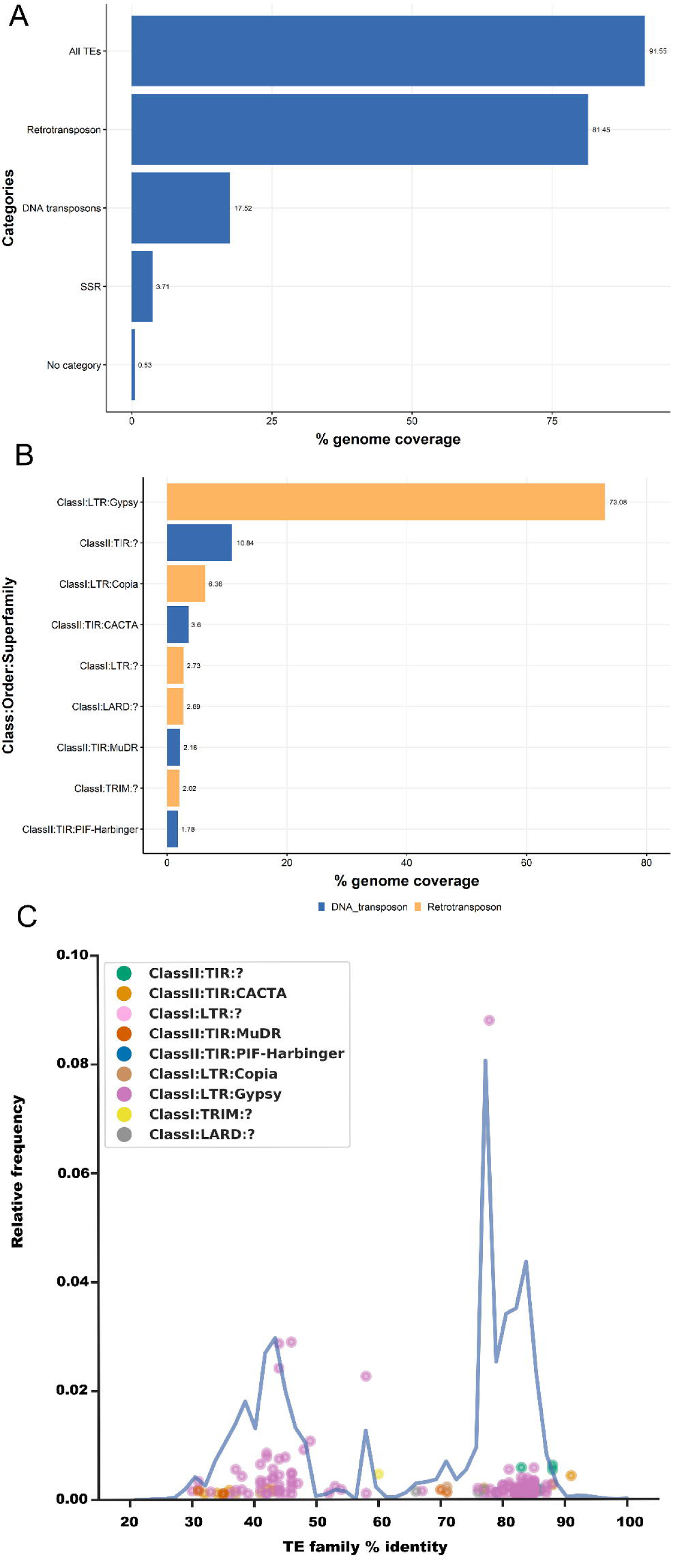
Repetitive element annotation on primary scaffolds. *Austropuccinia psidii* has a high repeat content driven by expansion of the Gypsy superfamily. Repetitive element annotation on primary scaffolds. (A) Percentages of genome coverage for all repetitive elements and different subcategories with some overlaps. These include transposable elements (TEs) of class I (RNA retrotransposons) and class II (DNA transposons), simple sequence repeats (SSR), and unclassifiable repeats (no category). (B) Percentages of genome coverage of Class I and Class II TEs categorized to the class, order, and superfamily levels wherever possible. Repetitive elements were identified using the REPET pipeline, and classifications were inferred from the closest BLAST hit (see Materials and Methods). The TE order is color coded in each Class I and Class II TE plot. (C) A subset of transposable element (TE) superfamilies has driven the genome expansion of *A. psidii*. The blue line indicates the mean TE family percentage identity distribution relative to the consensus sequence of TE families as a proxy of TE age. Individual points indicate the relative frequency of a specific TE family plotted at their mean percentage identity relative to the consensus sequence. Points are color coded according to the TE superfamily. Only highly abundant TE families are included in the plot.

### AT-rich transposable elements are likely due to 5-methyl-cytosine to thymine transitions

The high percentage of TEs and the overall low GC content prompted us to investigate the presence and distribution of AT-rich regions in the *A. psidii* genome. We identified AT-rich regions using OcculterCut (Testa et al. 2016), a program specifically designed to identify AT-isochores. We find two peaks with the major peak identified at around 33% GC content and a second smaller peak at around 41% GC content (Figure 2 A). The major peak largely overlaps with the GC content profile of TEs, while the second peak overlaps with the GC content of genes in the *A. psidii* genome. We investigated if these AT-rich regions are specific to *A. psidii* or if they are present in other rust fungi. All other analyzed rust fungi showed a single peak in the GC profile analyzed with OcculterCut (Additional Figure 1). For example, *P. striiformis* f. sp. *tritici* displayed only a single peak at around 45% GC content (Figure 2 B and Additional Figure 1). We further investigated the GC-content for two scaffolds that have telomeric start and end regions indicating putative pseudochromosomes, APSI_P025 (8 512 731 bp) and APSI_P027 (3 316 254 bp). Plotting of gene density and repeat density showed a lack of repeat or gene-rich islands, and instead an accumulation of TEs between genes (Figure 2C). Also, the AT-rich regions identified with OcculterCut did not correlate clearly with any specific genome feature.

Repeat induced point mutations (RIP) have been shown to contribute to AT-rich isochores in many ascomycetes (Gladyshev 2017). This process is commonly not found in basidiomycete fungi, yet several recent studies suggested RIP or RIP-like processes are present in Pucciniomycetes, including rust fungi (Horns et al. 2012; Amselem et al. 2015). We searched for the two DNA methyltransferases known to be involved in RIP in *Ascobolus immersus* and *Neurospora crassa* (Gladyshev 2017; Möller & Stukenbrock 2017) in our *A. psidii* proteome. We found clear homologs for both in addition to other DNA methylation related enzymes (Additional Table 1B). This suggests that the 5mC and potentially the 6mA DNA methylation pathways are intact in *A. psidii*, similar to a recent report in *P. graminis tritici* (Sperschneider et al. 2020). We calculated two dinucleotide frequency indices TpA/ApT and (CpA + TpG)/(ApC + GpT) to test for the presence of RIP. These have been used previously in *N. crassa*, *Rhynchosporium commune*, and *Marssonina brunnea* to confirm RIP and in *Parauncinula polyspora* and *Blumeria graminis* f. sp. *hordei* to exclude RIP (Margolin et al. 1998; Frantzeskakis et al. 2019) (Figure 3 A and B). We did not observe any indication of classic RIP in the *A. psidii* genome using these two indices because the two most abundant TE superfamilies (ClassI:LTR:Gypsy and ClassII:LTR:Copia) did not show any alterations in these indices when compared to the non-TE genome background. This contrasted with the genomes of *R. commune* and *M. brunnea* used as positive controls for RIP (Figure 3 A and B).

DNA methylation is directly linked to the silencing of transposable elements in plants and fungi (Wyler et al. 2020; Sperschneider et al. 2020). DNA methylation of 5mC sites was reported for *P. graminis* f. sp. *tritici* (Sperschneider et al. 2020) with a strong preference for CpG dinucleotide context. *Austropuccinia psidii* encodes all the required DNA methylation machinery (Additional Table 1B) and hence we hypothesize that DNA methylation is present and active in

*A. psidii*. Methylation of cytosine in the 5mC context is known to be hypermutable due to frequently occurring deamination of the methylated cytosine leading to transitions to thymine. Over time the subsequent G-T mismatch repair is imperfect and leads to an increased AT content in originally hypermethylated regions, including TEs (Maumus & Quesneville 2014). We tested whether a similar process occurs in *A. psidii* by analyzing the GC-content of TE families according to their age measured using family level identity as a proxy for age. Indeed, we found that the consensus sequence younger TE families (> 90% family level identity) displayed a higher GC content, and that this GC content decreased with TE family age of older TE families (Figure 3 C). This increase in AT-richness correlated with the depletion of CpG dinucleotides in individual TE copies over time (Figure 3 D). This finding suggests that the low GC-content of *A. psidii* is a consequence of rapid historical TE expansion followed by deamination of hypermethylated TEs and imperfect G-T mismatch repair.

**Fig. 2.**
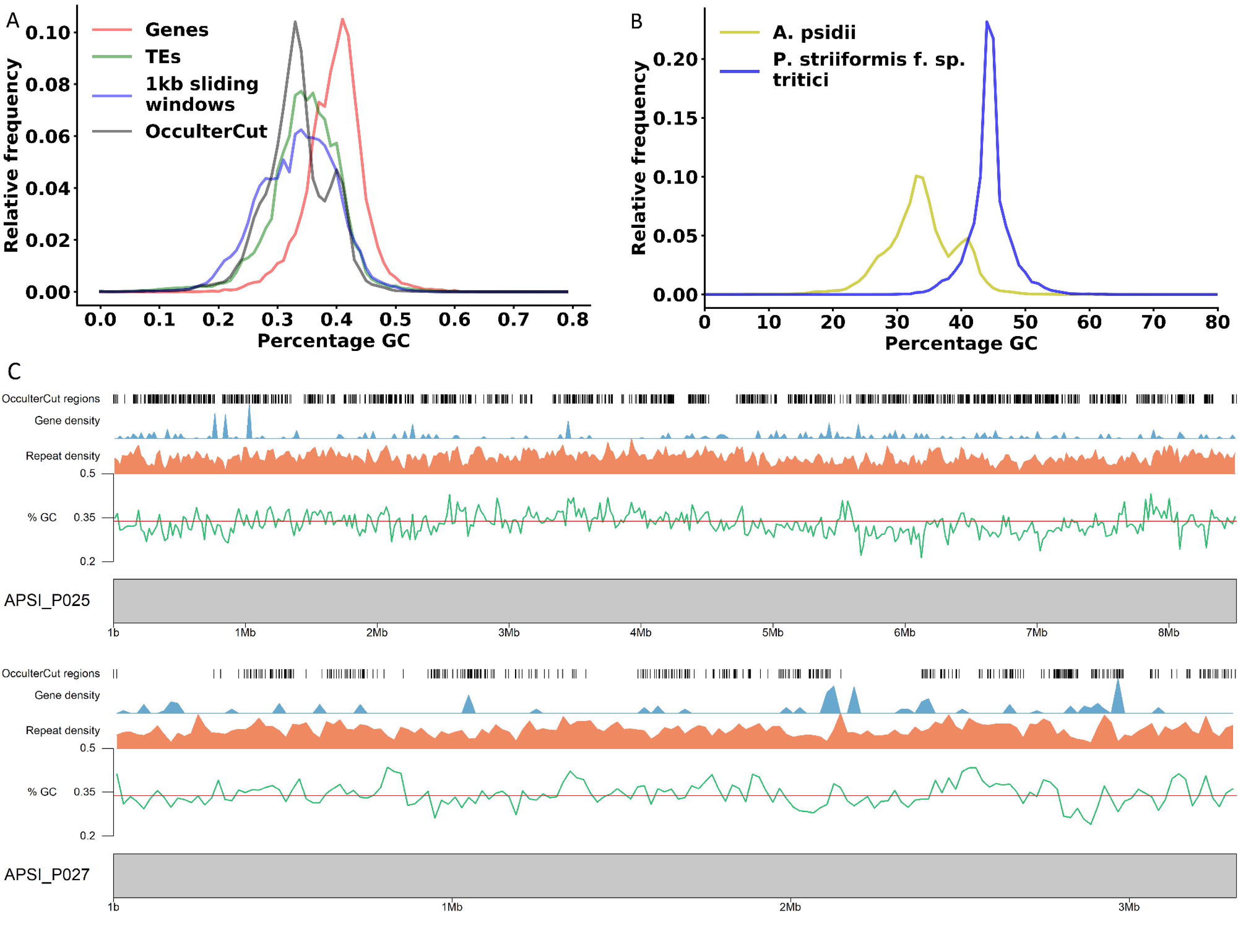
AT-richness in the *Austropuccinia psidii* genome is caused by transposable elements (A) *A.psidii* displays two distinct genome compartments in relation to GC content. Relative GC content of genome regions identified by genes, transposable elements (TEs), 1 kb sliding windows, or as identified by OcculterCut are shown. (B) AT-enriched regions are specific to *A. psidii.* Relative GC content of genome regions identified by OcculterCut in *A. psidii* and *Puccinia striiformis* f. sp*. tritici* (see also Additional Figure 1). (C) Karyoplots of scaffolds APSI_025 and APSI_P027. Gene and repeat density are shown at 20,000 bp windows. Mean GC-content of 33.8% is shown with a red line. OcculterCut regions of GC-content segmentation are shown as black lines.

**Fig. 3.**
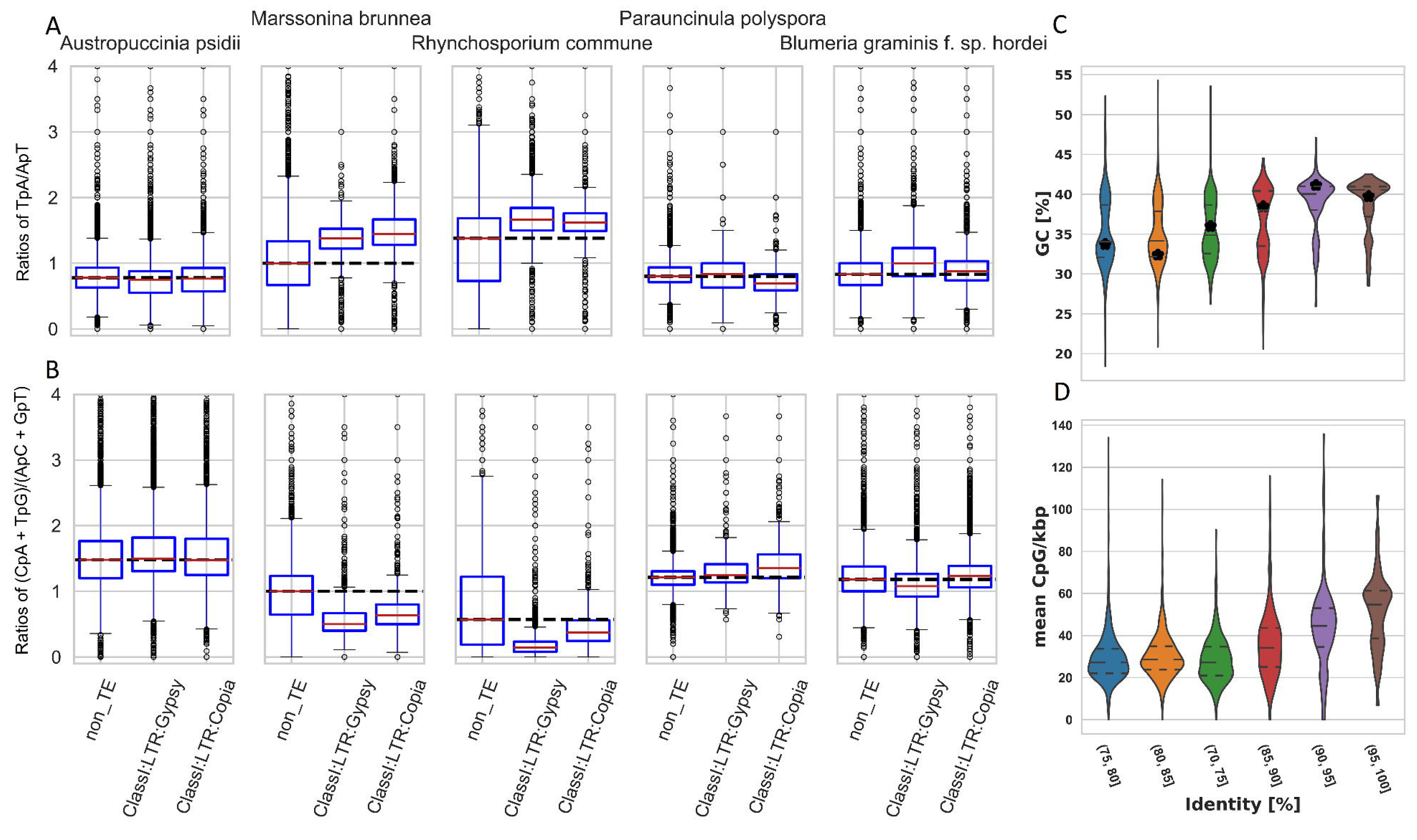
Depletion of CpG dinucleotides in transposable elements leads to AT-richness over time and not classic RIP mutations. *A. psidii* does not show classic RIP signatures in its TEs (A, B). (A) TpA/ApT and (B) (CpA + TpG)/(ApC + GpT) ratios in *A. psidii* and two ascomycetes with and without RIP mutations. Dinucleotide ratios are plotted for regions grouped according to their identity of non-TE, Gypsy, and Copia superfamily. The horizontal dashed black line indicates the median of the dinucleotide ratio in non-TE regions. Deviation from this median in other genome regions indicates RIP mutations as shown in *Marssonina brunnea* and *Rhynchosporium commune*. AT-richness and depletion of CpG increase over time in TE sequences of the *A. psidii* genome (C, D). (C) Percentage GC content of TE consensus sequences grouped by percentage identity relative to the consensus sequence as proxy for TE age. Plot shows younger TEs on the right. Horizontal lines show median and quantiles. Black stars indicate the weighted mean relative to genome coverage. Kruskal-Wallis H-test indicates a significant difference between the samples (p-value < 2.1e-57). Lines with * indicate significant differences with p-values < 1e-5 using Mann-Whitney U Tests with multiple testing correction. (D) Mean CpG content per kbp of individual TE insertions grouped by percentage identity relative to the consensus sequence as proxy for TE age. Plot shows younger TEs on the right. Horizontal lines show median and quantiles. Kruskal-Wallis H-test indicates a significant difference between the samples (p-value = 0.0).

### *Austropuccinia psidii* has extraordinarily long telomeres

We identified unusually long telomeres in the *A. psidii* genome and transcripts showed the presence of the predicted telomerase gene, indicating a putative role in maintaining telomeres during active growth. Prior to scaffolding the assembled contigs, we identified 29 telomeric regions at the start or end of primary contigs and 23 in the secondary contigs, based on the hexamer TTAGGG(n) (Lucía et al. 2010). We used these numbers to guide scaffolding and checked the telomere locations post scaffolding. Three primary scaffolds; APSI_P012 (37 965 047 bp), APSI_P025 (8 512 731 bp), APSI_P027 (3 316 254 bp), have telomeric regions at both the start and end indicating potential complete chromosomes. The final hexamer repeat numbers and lengths (Table 2) indicate extraordinarily long telomeres, over 1000 bp in length for 24 of the scaffolds, when compared to other fungi which have telomeres in the size range of 100-300 bp (Lucía et al. 2010; Pérez et al. 2009; Schwessinger et al. 2020; Sperschneider et al. 2020). Telomerase related transcripts were expressed, including telomerase reverse transcriptase (RNA-dependent DNA polymerase) and telomerase ribonucleoprotein complex -RNA binding domains (PF12009 and PF00078 protein ID: APSI_P017.12297.t3 and APSI_H002.12793.t2).

**Table 2.** Telomere repeat regions identified for the *Austropuccinia psidii* primary assembly before and after scaffolding.

| n(TTAGGG) | >1000 | 800-999 | 500-799 | 90-499 | <25 | Total |
| --- | --- | --- | --- | --- | --- | --- |
| Av. length (nt) | 8 159 | 5 223 | 3 754 | 1 409 | 87 |  |
| Pre-scaffold | 6 | 6 | 4 | 8 | 5 | 29 |
| Post-scaffold | 10 | 3 | 2 | 9 | 5 | 29 |

Numbers and average length in nucleotides (nt) of contigs/scaffolds with n(TTAGGG) sequences. n = number of hexamers, for example >1000 means more than 1000 x (TTAGGG) or more than 6000 nt.

### *Austropuccinia psidii* encodes a proteome closely related to other rust fungi despite increased intergenic distances

Based on *in planta* RNA-seq data as supporting evidence, 18,875 and 15,684 protein coding genes were predicted within the primary and secondary assemblies respectively. Comparison of intergenic distance between *A. psidii* and other fungi within the order Pucciniales shows large expansions of intergenic distances in *A. psidii*, while gene lengths incorporating untranslated regions (UTRs) are similar across six species of the Pucciniales (Figure 4).

**Fig. 4.**
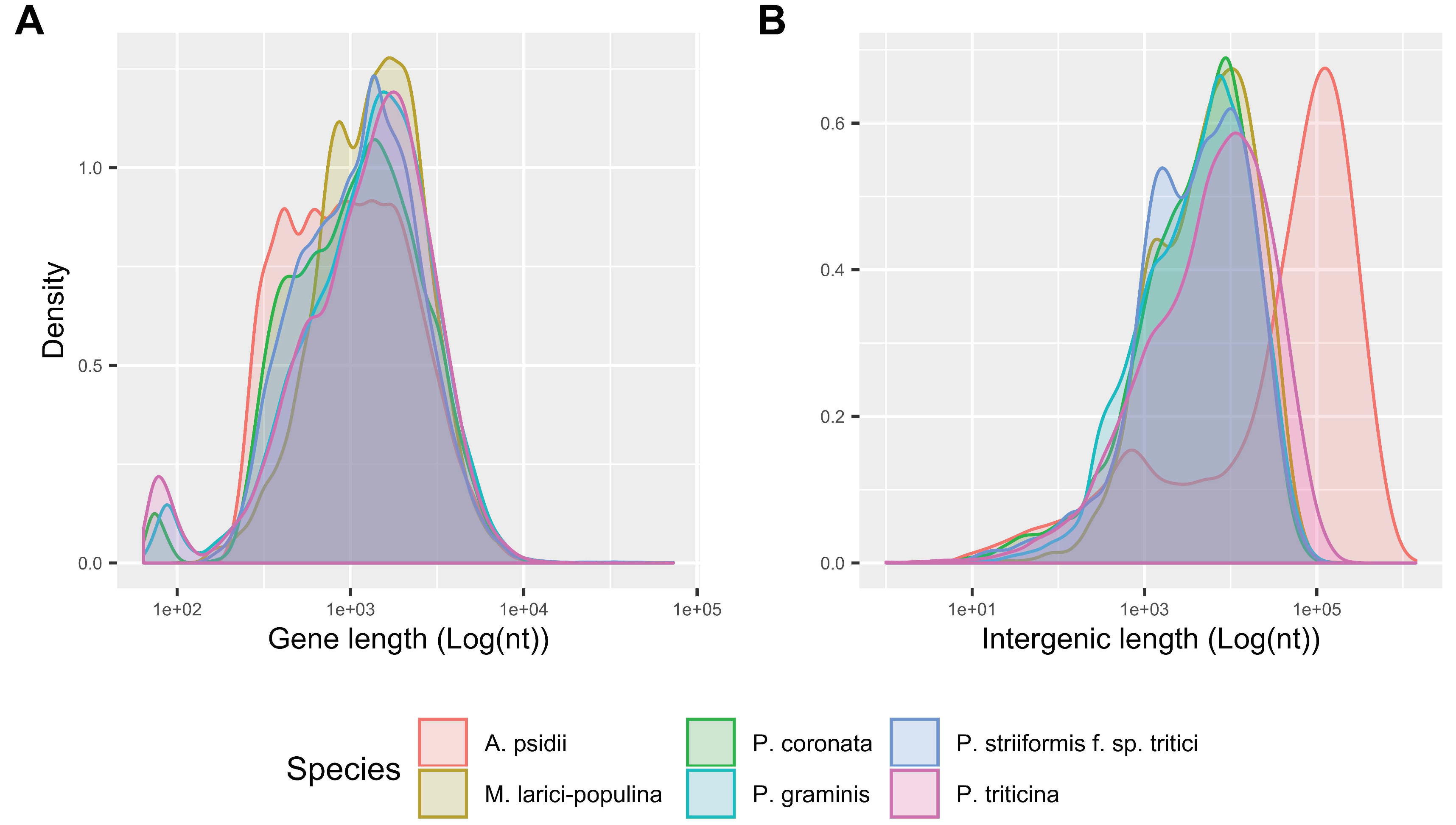
Structural annotation comparisons of gene (A) and intergenic (B) length, incorporating untranslated regions, across six species of Pucciniales reveal dramatically large intergenic expansions in *A. psidii*.

Putative effector genes were identified within the primary and secondary assemblies at 367 and 304, respectively. These putative effectors did not appear to be compartmentalized and had similar extended intergenic distances when compared to BUSCOs or all genes (Figure 5). Previous analyses have determined that apoplastic effectors are highly enriched in rust pathogens. On average 52% of effectors are localised to the apoplast compared to other fungal pathogens, around 25% (Sperschneider, Dodds, Singh, et al. 2018). We therefore predicted potential apoplastic proteins and effectors with ApoplastP (Sperschneider, Dodds, Singh, et al. 2018). We identified that 47% and 40% of predicted effectors are localized to the apoplast in the primary and secondary *A. psidii* assemblies respectively, indicating a potential non-cytoplasmic role in host manipulation. Expression validation, based on mapping *in planta* RNA-seq reads to the primary and secondary assemblies, was found for 3,923 and 2,435 sequences, including 78 and 63 predicted effectors. Overall, this suggests that *A. psidii* has a similar proteome size in terms of numbers when compared to other rust fungi (Duplessis et al. 2011; Schwessinger et al. 2018) (Table 3) and that TE insertions led to an overall increase of intergenic distances for most genes.

**Fig.5.**
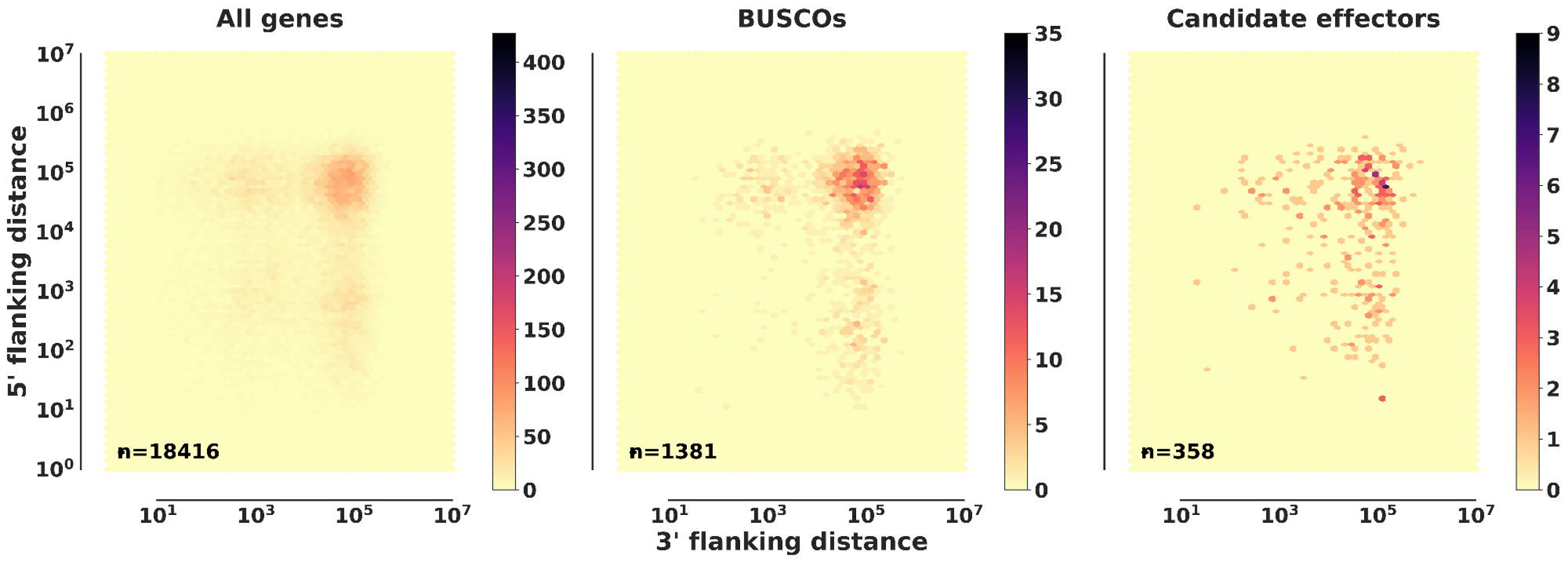
*Austropuccinia psidii* putative effectors are not found in gene sparse regions. Nearest-neighbor gene distance density hexplots for three gene categories including all genes, BUSCOs, and candidate effectors. Each subplot represents a distance density hexplot with the log10 3′-flanking and 5′-flanking distance to the nearest-neighboring gene plotted along the x axis and y axis, respectively.

To test overall proteome conservation between the related rust species we performed a comparative proteome analysis of the aforementioned rust fungi within the order *Pucciniales* including *A. psidii* using Orthofinder (Emms & Kelly 2019). We included proteomes from two tree rust species, *Cronartium quercuum* causing white pine blister rust (Pendleton et al. 2014) and *Melampsora larici-populina* causing poplar rust (Duplessis et al. 2011), to test whether protein orthologues may relate to life history of the host. Based on a species rooted phylogenetic tree from the multiple sequence alignment of single copy orthologues, *A. psidii* was placed closer to the cereal rusts than to the tree rust species (Figure 6 A). This finding is in accordance with previous analyses that examined the taxonomic placement based on ribosomal DNA and cytochrome c oxidase subunit 3 (COX3) of mitochondrial DNA of the type specimen for *A. psidii* (Beenken 2017; Aime et al. 2018). A graphic presentation of the orthologues shows shared ortholo-groups between *Pucciniales* species (Figure 6 B) indicating proteome closeness. However, a group of 101 orthologues (Figure 6 B, right) present in the tree rusts *M. larici populina*, *C. quercuum* and *A. psidii* appear to be absent from the cereal rust species (Additional Table 2).

**Fig.6.**
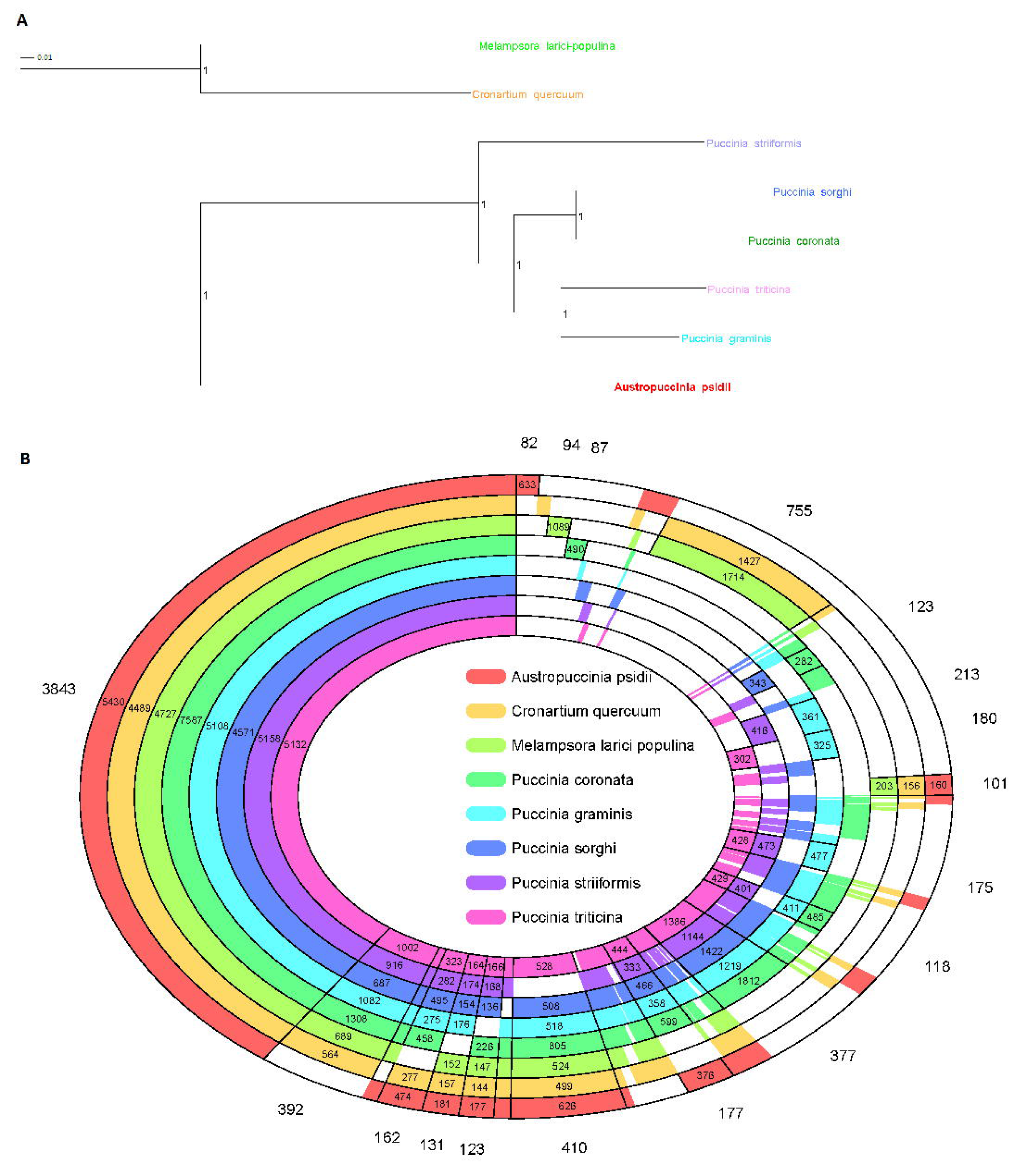
(A) Protein-protein species rooted tree based on multiple species alignment of orthogroups identified with Orthofinder (Emms & Kelly 2019). Scale represents substitutions per site and internal node values are species tree inference from all genes (STAG) supports (Emms & Kelly 2018). (B) Protein-protein comparisons across rust fungal species with each concentric ring indicating a different species. Numbers external to the rings represent counts of orthologue groups and numbers within each concentric ring represent the number of orthologue genes in that species per section. The figure shows every possible combination of species included in this proteome orthologue analysis, using concentric circles graphically present an overview of ‘closeness’ between the genomes. The species color code is consistent for Figures 5 and 6.

**Table 3.** Comparative assembly data across several available Pucciniales genomes. Abbreviated names indicated; A.p. (*Austropuccinia psidii* pandemic biotype from the current study in bold), P.s.t. (*Puccinia striiformis* f. sp. *tritici*), P.t. (*P. triticina*), P.g. (*P. graminis* f. sp. *tritici*), P.c. (*P. coronata*), P.s. (*P. sorghi*), M.l-p. (*Melampsora larici-populina*). *Predicted coding genes. nd=no data found.

| Common name (rust) | A.p. | P.s.t. | P.t. | P.g. | P.c. | P.s. | M.l-p. |
| --- | --- | --- | --- | --- | --- | --- | --- |
| NCBI/ENA | **PRJEB34084** | PRJNA396589 | PRJNA36323 | PRJNA18535 | PRJNA398546 | PRJNA277993 | PRJNA242542 |
| project |  |  |  |  |  |  |  |
| Genome (Mbp) | **1,018** | 83 | 135 | 89 | 150 | 100 | 101 |
| Scaffolds | **66** | 156 | 14,818 | 393 | 1,636 | 15,715 | 462 |
| N50 | **56 243 252** | 1 304 018 | 10 369 | 39 493 | 163 229 | 19 078 | 112 314 |
| L50 | **7** | 57 | 2 866 | 557 | 241 | 1 530 | 265 |
| Repeats (%) | **89** | 54 | 40 | nd | nd | nd | 45 |
| Coding genes* | **18 875** | 15 928 | 15 685 | 15 979 | 26 323 | 21 078 | 16 372 |
| GC (%) | **34** | 44 | 37 | 44 | 45 | 45 | 41 |

## Discussion

*Austropuccinia psidii,* the causal agent of the pandemic myrtle rust disease, is a dangerous and rapidly spreading plant pathogen. It infects perennial hosts in the family Myrtaceae, including iconic trees such as the eucalypts, and has an expanding global host list (Soewarto et al. 2019). Here we present the first highly contiguous genome assembly for *A. psidii* using DNA from an Australian isolate of the pandemic biotype that was previously assembled inadequately with short-read sequence data (Tan et al. 2014). We assembled the genome using long read sequence data to produce the largest assembled fungal genome to date at 1.02 Gbp haploid size.

The *A. psidii* genome is particularly large compared with the average fungal genome, around 44.2 Mbp (Tavares et al. 2014), and relative to its Myrtaceae hosts, around 300 - 650 Mbp (Thrimawithana et al. 2019; Myburg et al. 2014). Data based on flow cytometry determined the largest fungal genome at ∼3.6 Gbp (Egertová & Sochor 2017) and the largest known rust-type (Basidiomycota, Pucciniales) fungal genomes; *Gymnosporangium confusum* and *Uromyces bidentis*, at 893.2 Mbp and 2,489 Mbp, respectively (Ramos et al. 2015). Rust genomes appear to be larger on average than other fungal genomes and rusts infecting Poaceae (monocotyledon) are significantly smaller than those infecting Fabaceae (eudicotyledon) hosts, with average sizes of 170.6 Mbp, and 556.6 Mbp respectively. Interestingly, genome size does not appear to correlate with the perennial nature of host plants with the assembled genome of two important tree rust pathogens, *Melampsora larici-populina* (101 Mbp) (Duplessis et al. 2011) and *Cronartium quercuum* (22 Mbp) having modest-sized genomes, though sharing orthologs with *A. psidii* (Figure 6 B, right) that are absent in the cereal-infecting rust. While ploidy has been postulated as a possible reason for large fungal genome sizes (Egertová & Sochor 2017), the evidence to date is that genome expansion in rust species is due to repetitive elements, notably a proliferation of transposable elements (TEs) (Duplessis et al. 2011; Schwessinger et al. 2018; Foulongne-Oriol et al. 2013).

### Transposable elements shape the *A. psidii* genome

The large genome size in *A. psidii* is explained by the significant historical expansion of TEs with over 90% of the genome being made up of repetitive sequences. It is well known that TEs shape genome evolution and adaptation in animals (Kazazian 2004), plants (Bennetzen & Wang 2014), and fungi (Möller & Stukenbrock 2017). In fungi TEs have been implicated in genome compartmentalization in a subset of plant pathogens (Frantzeskakis et al. 2020), regulation of gene expression (Castanera et al. 2016), and evolution of avirulence effectors (Zhong et al. 2017; Salcedo et al. 2017). In addition, TEs themselves appear to be regulated by stress conditions (Lanciano & Mirouze 2018; Muszewska et al. 2019) such as during plant infection in the case of *Zymoseptoria tritici* (Fouché et al. 2020). This stress dependent TE expression might explain why many plants and fungi have highly variable TE insertion site landscapes at the population level were many loci are unfixed, as shown for *Arabidopsis thaliana* (L.) Heynh (Stuart et al. 2016; Quadrana et al. 2016) and *Z. tritici* (Lorrain et al. 2020; Oggenfuss et al. 2020). The TE variation within a species extends beyond variations of insertions sites. In *P. striiformis* f. sp. *tritici* TEs are highly enriched in presence-absence polymorphisms between high quality genomes of two major lineages (Schwessinger et al. 2020). In *Z. tritici* the TE landscape varies between different isolates in terms of TE content and TE insertion ages (Lorrain et al. 2020). Similarly, the comparison of four high quality maize genomes revealed extensive polymorphism at the family and individual TE level (Anderson et al. 2019). In the case of *A. psidii* TEs appear to be ‘species specific’ as we could not identify substantial TE insertions of *A. psidii* TEs in other Pucciniales even though the overall gene content was highly conserved within this order. These general observations led us to suggest the following hypotheses about *A. psidii*’s genome evolution.

First, we propose that *A. psidii* evolved from a small subpopulation of its predecessor with a random assortment of TEs. Within this population, the initial activation of TEs might have been beneficial to host adaptation and host range expansion as suggested by the “TE thrust hypothesis” (Oliver & Greene 2012). This hypothesis suggests that the time constrained activation of TEs generates genetic diversity that is useful for adaptation and contributes to speciation. In *A. psidii* we see a clear TE burst in recent history (>75% family level identity) which contributed significantly to its genome size expansion. This TE burst might coincide with the time estimate of *A. psidii*’s last common ancestor with *Dasyospora* spp., about 40 Mya years ago (Aime et al. 2018), and supports the finding that host jumps and host range expansion drive diversity in the Pucciniales (Mctaggart et al. 2015). Interestingly, this postulated split follows major speciation and radiation events in Myrtaceae starting about 60 Mya (Berger et al. 2016; Thornhill et al. 2019). Due to the small subpopulation size, highly expressed TEs would be fixed rapidly in the population and could contribute further to, or drive, reproductive isolation. These carrier subpopulation hypothesis effects (Jurka et al. 2011), might be exacerbated by long- periods of asexual reproduction in the *A. psidii* dikaryotic stage which does not allow for TE purging via meiotic recombination. Indeed, it has been recently suggested that the host of the dikaryotic urediniospores (telial host) is the major driver of biogenic radiation in rust fungi (Aime et al. 2018). Overall, the specific transposon composition of *A. psidii* might be the result of genetic drift and population bottlenecks and the mechanisms of genome expansions via the TE burst might be intrinsically linked to adaptation and host range expansion. In future, it will be interesting to investigate the population level variation of TEs in *A. psidii* and compare this with other rust fungi of similar genome size. As more high-quality contiguous rust genomes become available, it will be intriguing to explore the impact of past TE bursts on gene synteny in the Pucciniales and test if it follows the general phylogeny and is potentially involved in driving speciation and host range expansion.

Secondly, TE expansion in *A. psidii*, has led to an overall dramatic increase of intergenic distance, an order of magnitude greater when compared to six other Pucciniales species. This overall increase in intergenic distance suggests that some TE families insert preferentially into intergenic regions, including promoter sequences, as opposed to forming large gene poor TE rich islands. We did not observe such regions in *A. psidii*. This type of preferential TE insertion has been observed for certain Gypsy elements that target transcription initiation sites of RNA polymerase III-transcript genes (Jurka et al. 2011). It has been determined generally that the larger the fungal genome, with greater distances between genes, the fewer TEs overlap with genes (Muszewska et al. 2019). Future studies might address whether Gypsy families, which contributed to the genome expansion in *A. psidii*, show an insertion site preference for initiation sites and if this correlates with the chromatin accessibility regulated by DNA and histone modifications. This preferential TE insertion site profile might also explain the absence of any kind of ‘two-speed genome’ type of compartmentalisation in rust fungi (Frantzeskakis et al. 2020).

### *Austropuccinia psidii* AT richness is a direct consequence of transposable element silencing

In a population context the purging of TEs from a genome is competing with the TE insertion site preference when defining the TE insertion site spectrum across a population. The purging of TEs is linked to recombination during meiosis and sexual reproduction and in the case of extended asexual reproduction cycles, this mechanism of genome defence against TEs is limited. Another genome defence mechanism against TEs is the methylation of TEs which initially leads to TE silencing (Deniz et al. 2019). This initial silencing is often followed by continuous deterioration due to accumulation of mutations via deamination of 5mC causing transitions from cytosines to thiamines. Methylated 5mC is more liable to spontaneous deamination than unmethylated cytosines (Walsh & Xu 2006) and the resulting T:G mismatch repair is less efficiently repaired than U:G mismatch repairs arising from spontaneous deamination of non-modified cytosines (Krokan & Bjoras 2013). These biochemical processes lead to the reduction of GC-content in heavily methylated TEs over time as shown in *A. thaliana* for example (Maumus & Quesneville 2014). We observed a similar pattern for TEs in *A. psidii* with older TE families having a lower GC content. This coincides with a depletion of CpG dinucleodites in older TEs. Overall, this is consistent with our recent findings that TEs are heavily methylated in

*A. P. graminis* f. sp. *tritici* and that 5mC methylation has a CpG dinucleotide context preference (Sperschneider et al. 2020). Our observation is also consistent with recent reports that suggest a preferential mutation dinucleotide context of CG in TE sequences of several fungal species (Amselem et al. 2015; Neafsey et al. 2010) and might also explain the ‘RIP-like’ signatures reported for some basidiomycetes including two Pucciniales (Horns et al. 2017, 2012). Hence, we suggest that the AT-richness in *A. psidii* is a simple consequence of biochemical instability of CpGs in TEs combined with insufficient DNA mismatch repair. A similar effect has been recently observed in *Z. tritici* with methylation mediated mutations in the CpA dinucleotide context. The loss of the de novo methyltransferase DIM2 leads to reduced methylation and reduced mutability of CpA, but not CpG, dinucleotides. Restoring the DIM2 in *dim2* mutants increases the overall mutation rate and most of the de novo mutations are in a CpA context (Möller et al. 2020). It is intriguing that the 5mC mediated mutation effect is the same overall but in different dinucleotide contexts when comparing some ascomycetes such as *Z. tritici*, *N. crassa*, with basidiomycetes, for example the rust fungi. The different dinucleotide preferences might be related to differences in efficiency of G:T mismatch repairs depending on the (di-) nucleotide context. It will be interesting to more widely explore the conservation of canonical DNA repair machinery proteins in fungi and the potential correlation with enhanced 5mC turnover rates (Bewick et al. 2019).

### Isolate variation is a concerning threat to native flora

A preliminary analysis of SNP data across Australian *A. psidii* isolates, using a Hawaiian pandemic isolate and a Brazilian biotype for comparison (Additional Data 1.docx), has raised some concerns regarding potential isolate variation within Australia. These results indicate that new incursions may have already occurred and/or that genetic variants are occurring in Australia. Changes in *A. psidii* virulence have been observed on species previously considered resistant or highly tolerant including *Acmena ingens* (F.Muell. ex C.Moore) Guymer & B.Hyland (syn. *Syzygium ingens*) and *Waterhousea floribunda* (F.Muell.) B.Hyland, with any symptoms observed restricted to limited leaf spots (Pegg et al. 2014). More recent surveys (G Pegg, personal communication, 25 November 2019) have identified significant levels of infection on both these species resulting in severe foliage blighting and dieback caused by *A. psidii* infection. In the case of *W. floribunda*, specific trees that have not previously shown infection, despite other species in the area recorded as infected back in 2012, have now succumbed to infection. Similarly, mature *Syzygium corynanthum* (F.Muell.) L.A.S. Johnson trees that were considered to have no or low levels of infection in 2016 (Pegg et al. 2017) now present extensive levels of dieback and are in severe decline. Several Australian plants that were previously wide-spread are now listed as threatened or critically endangered since the arrival of the pathogen in 2010 (Fernandez Winzer et al. 2019; Berthon et al. 2019; Makinson 2019) and similar concerning impacts have been noted in New Caledonia (Soewarto et al. 2018). The contiguity of this genome assembly will aid the identification and containment of newly introduced and potentially more virulent isolates and provides an excellent reference for the study of *A. psidii* evolution and adaptation. The isolation of the Australasian region from other parts of the world has provided rare opportunities to study long-term evolution in rust pathogens of annual cereal crops (Park et al. 1995), and similar opportunities now exist to study evolution in a rust pathogen of perennial hosts in native ecosystems.

### Potential implications of the research

We have generated a highly contiguous reference genome assembly for the pandemic biotype of *A. psidii* using long-read sequencing and chromosomal confirmation capture sequencing technologies. The resource of a genome assembly from long reads has enabled the spanning of highly repetitive sequence regions, previously problematic to assemble with short read sequence data. We used *in vivo* expression data for genome annotation and have identified genome features such as extended telomeres and very extensive repetitive regions dominated by transposable elements. At 1.02 Gbp this is the largest fungal genome yet to be assembled and provides a foundation for future molecular research as well as a base-line resource to monitor the spread of the pathogen. It is expected that this genome resource will enable improved biosecurity management by monitoring pathogen populations to detect new incursions and population shifts and permit a deeper understanding of the molecular interaction between the pathogen and the host plant.

## Methods

### Austropuccinia psidii spores

*Austropuccinia psidii* urediniospores were initially collected from the host plant *Agonis flexuosa* (willow myrtle) in 2011, Leonay, NSW, and a single pustule isolate was increased through five cycles on the susceptible host plant *Syzygium jambos* (rose-apple) (Sandhu et al. 2015). Resulting spores (isolate ID: Au_3, culture no. 622 and accession 115012) were maintained in liquid nitrogen at the Plant Breeding Institute, (PBI) University of Sydney and used for all subsequent workflows including flow cytometry, Hi-C libraries and high molecular weight (HMW) DNA extraction.

### Nuclei size determination using flow cytometry

We estimated nuclear DNA content of *A. psidii* cells using flow cytometry. The fungal nuclei fluorescence intensity were compared with that of *Eucalyptus grandis*, as an internal standard, 2C=1.33pg, 640 Mbp (Grattapaglia & Bradshaw Jr 1994; Myburg et al. 2014). Nuclei were extracted from cells using methods previously described (Tavares et al. 2014) with slight modifications. Approximately 50 mg of leaf material from *E. grandis* (internal standard) were chopped in a petri dish with a sterile blade in 1 mL of chilled, modified Wood Plant Buffer (WPB) (Tavares et al. 2014) consisting of 0.2M Tris-HCl, 4 mM MgCl_2_, 0.1% Triton X-100, 2 mM Na_2_EDTA, 86 mM NaCl, 20 mM Na_2_S_2_O_5_, 1% PVP-40, pH 7.5. New leaves were taken from pathogen free *E. grandis* plants and *E. grandis* plants with *A. psidii* infection symptoms one-week post-inoculation. Inoculated plants were treated using previously described methods (Tobias et al. 2018). Hence, the inoculated leaves were used to determine the *A. psidii* fluorescence peaks in relation to the un-inoculated leaves as the reference standard. RNase A (50 μg/mL) was added and samples were incubated on ice for 15 minutes prior to centrifugation through 25 μm nylon mesh. Two DNA stains were added to the samples; 4’,6-diamidino-2-phenylindole (DAPI) (5 μM) and propidium iodide (PI) (50 μg/mL) (Sigma-Aldrich, St. Louis, USA) and incubated at room temperature for 30-60 minutes. Fluorescence intensity was measured for 10, 000 events per sample, threshold rate of 370 events per second and flow rate medium, with BD FACSVerse™ using laser settings for PI and DAPI. A minimum of three replicates were run for the sample and the reference standard. Calculations were made using the reference standard (un-inoculated *E. grandis* leaves) compared to the inoculated sample. Histograms, using linear scale, were gated to reduce background fluorescence for each replicate and descriptive statistics obtained for the peaks using Flowing Software (v2.5.1) (Terho 2013). Medians with maximum coefficient of variation < 8.3 were determined. Calculations to determine genome size were based on taking the ratio of the median fungal fluorescence intensity (FI) divided by that of the internal standard, in this case uninfected leaves of *E. grandis* (Doležel & Bartoš 2005). A minimum of three replicates per sample were used. It was assumed that the dominant cell cycle stage (G1) FI peak for the rust fungus was the 1C content, due to the dikaryotic nuclei released from cells, whereas the dominant peak for the reference was 2C (Tavares et al. 2014). The nuclear DNA content in picograms (pg) was then multiplied by 978 Mbp (Doležel & Bartoš 2005) to estimate the genome size (1C) for the rust. Flow cytometry results indicated the 1C nuclei at 1.42 pg (1,388.91 Mbp).

### K-mer genome size estimation

As an additional approach to predict the genome size of *A. psidii*, raw Illumina data (SRR1873509) (Tan et al. 2014) was downloaded from the National Centre for Biotechnology Information (NCBI). Data was run through Jellyfish software (v.2.2.6) (Marçais & Kingsford 2011) with k-mer size of 16 and 21 and the following parameters: count -t 8 -C -m (16) 21 -s 5G. Histogram outputs (Jellyfish v.2.2.6) were visualised and coverage determined at 3X. The diploid genome size was calculated at 2,100 Mbp by dividing the total number of k-mers (21mer) by the mean coverage. From these results a haploid genome of around 1 Gb was predicted.

### DNA extraction and sequencing

For long read sequencing a modified CTAB HMW DNA extraction procedure (Schwessinger 2016) was used with the modification of using RNase A (Qiagen, Australia) instead of RNase T1. One gram of frozen *A. psidii* spores (isolate ID: Au_3, culture no. 622 and accession 115012) were ground in liquid nitrogen and then all methods were run up to step 33 of the protocol (Dong 2017). The DNA solution was then further purified to separate HMW DNA from other impurities and low molecular weight DNA as described in the protocol. DNA concentration and purity were measured with a Qubit 3 (Invitrogen) and Nanodrop ND-1000 (Thermo Fisher Scientific). If the ratio of concentration obtained from the Qubit to that of Nanodrop was smaller than 0.5, AMPure beads (Beckman, Coulter Inc.) were used to purify the DNA at aratio of 0.45 beads to 1.0 DNA (vol/vol) following the manufacturer’s protocol. We obtained a final quantity of around 240 μg of DNA and 10-15 μg are required for one library. DNA integrity was checked by pulsed-field gel electrophoresis. The DNA was first sequenced on PacBio RSII (18 SMRT cells) at Ramaciotti Centre for Genomics, University of NSW, Australia. Later, samples were sequenced on PacBio Sequel (total 11 SMRT cells). The sequencing library was prepared using SMRT cell template prep Kit 1.0-SPv3 with BluePippin Size-selection with 15-20 kb cutoff, and then sequenced using Sequel Sequencing Kit 2.0 at the Plant Genomics Centre, School of Agriculture, Food and Wine, the University of Adelaide, and the Ramaciotti Centre for Genomics. Altogether, a total of 126.1 (bax.h5) and 110.5 (subreads.bam) Gigabytes (GB) of raw data was obtained. The data for this study are deposited at the European Nucleotide Archive (ENA) at EMBL-EBI under study accession: PRJEB34084 and sample accession ERS3953144.

### RNA isolation and sequencing

*Syzygium jambos* young leaves were infected with myrtle rust according to a previously determined high inoculum protocol (Tobias et al. 2018). Infected leaves were harvested at 6, 12, 24, 48 hours, 5, and 7 days post-inoculation and stored at -80 °C. Total RNA was extracted from about 100 mg tissue, using the Spectrum Total RNA Kit (Sigma-Aldrich, St. Louis, MO), protocol A. The total RNA was then treated with RNase-free DNase I (New England BioLabs Inc.), and column purified using ISOLATE II RNA Mini Kit (Bioline, Australia) according to the manufacturer’s instruction. The quantity and quality of the total RNA were examined by Nanodrop (Thermo Scientific) and Agilent 2100 Bioanalyzer (Agilent Technologies). Library was constructed using a TruSeq Stranded mRNA-seq preparation kit (Illumina Inc., San Diego, CA, USA) and sequenced on NextSeq 500 (PE 150bp) in one MID flowcell at Ramaciotti Centre for Genomics, University of NSW, Australia. Data thus derived were used to assist genome annotation.

### Genome assembly

We optimized the sequencing based on our predicted genome size of ∼1 Gbp and sequenced 29 SMRT cells (18X RSII plus 11X Sequel). After converting RSII data to subreads.bam we obtained 162 GB of data. Details on data pre-processing and assembly are available at the myrtle-rust-genome-assembly github repository (Tobias 2019). In brief, RSII bax.h5 files were converted to subreads.bam before fasta files were extracted from all datasets. After extracting fasta files for assembly we had approximately 72X raw sequence coverage (7.24E+10 bases and 5.80E+06 reads). The reads were then mapped to an in-house *A. psidii* mitochondrial sequence to retain only genomic DNA sequence data (APSI_mitochondria.fa available on the github repository) (Tobias 2019). Canu v1.6 long read assembly software (Koren et al. 2016) was used to assemble the genome with correctedErrorRate=0.040, batOptions="-dg 3 -db 3 -dr 1 -ca 500 -cp 50" and predicted genome size of 1 Gb. The assembly ran for over four months continually on the University of Sydney High Performance Computing (HPC) cluster. Approximately 359,878 CPU hours were used for the assembly stage. Grid options were specified, and job scheduling was managed with PBS-Pro v13.1.0.160576, thereby limiting the number of jobs running at any one time to around 400. Each job required 4 CPUs, around 16 GB memory allocation and progressive allocations of walltime to a maximum of 70 hours. The primary genome assembly file, named APSI_primary.v1.fasta, has been deposited under the following ENA accession: ERZ1194194 (GCA_902702905). Locus tags are registered as APSI (for *<u>A</u>*ustropuccinia *<u>psi</u>dii*) and scaffolds identified as APSI_Pxxx (where x indicates scaffold number) for primary assembly. The diploid assembly file (APSI_v1.fa) is available from DOI:10.5281/zenodo.3567172 and incorporates the 67 secondary assembly scaffolds (APSI_Hxxx for secondary/haplotigs).

### Deduplication of assembly and polish

To deduplicate the assembly into primary contigs and secondary haplotigs we used the Purge Haplotigs (v1.0) pipeline (Roach et al. 2018) with alignment coverage of 65%. We polished the primary assembly using the genomic consensus algorithm *arrow* within Anaconda2 (v.5.0.1) (PacBio), after aligning with Minimap2 (v2.3) (Li 2018) and processing the data for compatibility, as described (Tobias 2019). This process was repeated twice and genome completeness for the primary and secondary genomes was assessed with BUSCO (v3.1.0) (Simão et al. 2015) using genome mode and the Basidiomycota database (basidiomycota_odb9 downloaded 24/02/2017) of 1,335 conserved genes. Where conserved single copy orthologues were present on the secondary contigs and absent from the primary contigs, these contigs were detected and moved to the primary assembly to ensure inclusion of conserved genes in downstream analyses. Taxonomy classification based on final assemblies was carried out with Kraken2 (Wood & Salzberg 2014) against NCBI/RefSeq Fungi database (dated 20190905) and visualized with Krona (v.2.7) (Ondov et al. 2011) (Additional Figure 2.HTML).

### Telomere identification for scaffolding

Prior to scaffolding we first checked for telomeric regions using blast+ v.2.3.0 (blastn e-value 1e-5) (Altschul et al. 1990) based on the hexamer TTAGGG(n) (Lucía et al. 2010) with the aim of guiding the scaffold numbers to putative chromosomes. We tested n=20, 40, 80, 120. From n=40 (length=240 bp) the contigs and numbers remained consistent with 100% match and e-value=0 for 24 contigs. A further five contigs had e-values less than 1.00E-07. Further analysis with seqkit (v0.10.1) (Shen et al. 2016) operation *locate* and a minimum TTAGGG repeat of 8 (and then 6) confirmed the BLAST findings. The hexamer repeat locations were at the start and/or end of contigs and were visualized using Integrated Genomics Viewer (IGV v2.6.3) (Thorvaldsdóttir et al. 2013). We reasoned that the repeats were likely to represent telomeric regions and used these numbers to guide Hi-C scaffolding.

### Hi-C libraries and scaffolding

Hi-C libraries were prepared using the Phase Genomics, Inc. (www.phasegenomics.com) Proximo Hi-C Kit (Microbe) to the manufacturer specifications. Illumina paired-end reads (125 bp on HiSeq) generated from Hi-C libraries were used to scaffold contigs. Reads were firstly trimmed with adapters and low-quality bases, and then mapped to the primary and secondary assemblies independently following the mapping workflow from Arima Genomics (2019). Three Hi-C scaffolding programs were tested (LACHESIS, SALSA and ALLHIC) before settling with ALLHIC (v.0.8.11) (Zhang et al. 2019). We tested the grouping, ordering and orienting of the contigs with both MluCl and Sau3AI, (SALSA) as enzyme cutting sites, both together and separately (LACHESIS and ALLHIC). Final scaffolds were generated from ALLHIC with the use of MluCl (“AATT”) as the cutting site.

### Transposable elements (TE) analysis

Repeat regions were annotated as described previously (Schwessinger et al. 2018) using the REPET pipeline (v2.5) for repeat annotation in combination with Repbase (v21.05). For de novo identification, we predicted repeat regions with TEdenovo in the initial assembly by subsampling 300 Mb of sequence space at random to reduce compute time with the risk of losing low abundant TEs. We called the resulting repeat library MR_P2A_300Mb. We annotated the primary assembly with three repeat libraries (repbase2005_aaSeq, repbase2005_ntSeq, and MR_P2A_300Mb) using TEanno from the REPET pipeline. Detailed description of the repeat annotation and analysis can be found in the associated github repository https://github.com/Team-Schwessinger/TE_myrtle_rust. Most of the RIP index analysis, GC-content and CG:GC depletion analysis of TEs was inspired by previous publications (Schwessinger et al. 2020). We identified AT-rich regions using OcculterCut (Testa et al. 2016). We calculated GC content for specific genome features or sliding windows (window size 1kb, slide 200bp) using bedtools (Quinlan & Hall 2010). We calculated dinucleotides using facount (Kent et al. 2002). Karyplots of scaffolds were plotted with karyplotR (Gel & Serra 2017).

### RNA data analysis

RNA-seq data were generated for six time points. Each set of data for different time point was processed in parallel in a consistent way. The quality of raw RNA-seq data was checked using FastQC (v.0.11.7) and an overall summary for all samples was created with MultiQC (v.1.5). Based on the QC reports, the data were cleaned using fastp (v.0.19.4) (Chen et al. 2018) trimming 15bp and 10bp from the 5’ and 3’ respectively, together with ‘--trim_poly_x --trim_poly_g’ trimming. The cleaned data for each time point were assembled independently using Trinityrnaseq (v2.6.5) (Haas et al. 2013). Six transcriptome assemblies were merged to create the final transcripts using Evigene (v.18-01-2018) (Gilbert 2013). Assembled transcripts were merged for a total of 80,804 representative genes and 108,659 alternative forms. Of the total transcripts, 8% and 9% were classified as fungi in the main and alternative gene sets, respectively (Additional Figure 2.HTML).

Taxonomy classification based on final assemblies, as well as representative and alternative forms of genes within transcriptomes, was subsequently carried out with Kraken2 (Wood & Salzberg 2014) against NCBI/RefSeq Fungi database (dated 20190905) and visualized with Krona (v.2.7) (Ondov et al. 2011) (Additional Figure 2.HTML). A large percentage of ‘no hit’ were recorded as plant genomes were not included in the database (NCBI/RefSeq Fungi). In addition to transcriptome assemblies, the cleaned data were mapped to both the *A. psidii* genome and the *Metrosideros polymorpha* (‘ōhi’a) genome, where the latter is the most closely related to the inoculated host plant among the publicly available ones. The *M. polymorpha* genome was downloaded from http://getentry.ddbj.nig.ac.jp and mapping was done with bowtie2 (v.2.3.4) (Langmead et al. 2009) in sensitive local mode. RNA-seq alignments showed that most reads were from the host plant and a minor percentage of data were from the fungal isolate. Mapping the RNA reads to the assembled *A. psidii* genome, the lowest overall mapping rate was observed in the sample collected at 6 h (0.77%) and the highest in 7 d (8.58%) post inoculation. Mapping to the *M. polymorpha* plant genome, the overall mapping rate was stable and greater than 76.7% in all samples except 7 d (which was 71.55%). Reads that did not map to either genome are likely to include mitochondrial reads and discrepancies with the plant host genome (genome of the species not currently available), thereby reducing mapping efficiency.

We mapped the combined, trimmed RNA-seq reads to the diploid assembly with STAR (v.2.7.2b) with the following parameters –sjdbGTFfile --sjdbGTFtagExonParentTranscript Parent --sjdbGTFtagExonParentGene ID --quantMode GeneCounts --outSAMtype BAM Unsorted --outSAMprimaryFlag AllBestScore. The output APSI_ReadsPerGene file was sorted by counts in descending order, per predicted coding regions, to identify sequences with expression. We took a cut-off of five mapped reads to one, and only one, predicted gene as expression evidence. When the cut-off was reduced to one mapped read, expression was found for 10,613 protein coding genes including 210 predicted effectors. The assembled transcriptome fasta file is available from DOI:10.5281/zenodo.3567172 and data (APSI_primary/secondary_v1.xlsx) are accessible from the github repository (Tobias 2019).

### Gene prediction and functional annotation

We annotated the primary and secondary scaffolds independently using Braker2 (Hoff et al. 2019) by first soft masking with RepeatModeler (v.1.0.8) (Smit & Hubley) and RepeatMasker (v. 4.0.6) (Smit et al. 1996). RNA-seq reads (29.4 GB) were trimmed with Fastp (v.0.19.6) (Chen et al. 2018), and mapped to the masked scaffolds with Hisat2 (Kim et al. 2015) for alignment files (RNA-seq hints). Structural annotation outputs from Braker2 were utilized in downstream functional annotation steps. Automated functional annotation was performed on the 18,875 (primary) and 15,684 (secondary) Braker2 predicted proteins for domain and motif searches with InterProScan (v.5.34-73.0) (Jones et al. 2014) (Additional Table 2). A BUSCO (Simão et al. 2015) analysis was run in protein mode against the combined predicted annotated genes with Basidiomycota database of 1,335 conserved genes. Additionally, we tested for transmembrane regions and signal peptide motifs using SignalP (v.4.1f) (Nielsen 2017). A reduced file of sequences conforming to these predictions was submitted to both EffectorP (v.2.0) (Sperschneider, Dodds, Gardiner, et al. 2018) and ApoplastP (v.1.01) (Sperschneider, Dodds, Singh, et al. 2018) to identify predicted effectors and apoplastic proteins from each scaffolded assembly. Predicted effector fasta files were submitted to the online HMMER (Potter et al. 2018), using phmmer and the reference proteomes database, for an alternative annotation output based on homology (Additional Table 3). In order to determine allelic counterparts within both the primary and secondary assemblies we ran blast+ (v.2.3.0) (blastp e-value 1e-5) (Altschul et al. 1990) with the predicted protein fasta files and identified query and subject equal aligned lengths at 100% match. Of the predicted effectors in the primary and secondary assemblies, exact amino acid duplicates were found for 97 sequences. Annotation files and multi-sequence fasta files for predicted coding sequences, amino acid sequences and effectors for the combined primary and secondary assemblies are available at DOI:10.5281/zenodo.3567172.

### Comparative analysis of Pucciniales proteins

We used Orthofinder v2.2.7 (Emms & Kelly 2019) with NCBI (DOE Joint Genome Institute) and Mycocosm (DOE Joint Genome Institute) (‘*’ represents data downloaded from Mycocosm) downloaded protein fasta files; \**Puccinia coronata* f. sp. *avenae*, *P. striiformis* 93-210, *P. graminis* f. sp. *tritici* CRL 75-36-700-3, *P. sorghi* RO10H11247, \**P. triticina* 1-1 BBBD Race 1, *Melampsora larici*-*populina* 98AG31, *Cronartium quercuum* f. sp. *fusiforme* G11. A visualized summary of orthologue clusters between species was generated with in-house PERL and R scripts. We visualized the evolutionary phylogeny, using the species rooted tree based on orthologs, with Dendroscope (v3) (Huson & Scornavacca 2012).

Additionally, we reviewed the literature to determine basic genome statistics for these rust pathogens and compared the datasets gene architecture based on annotations (gff/gff3 files). For the analysis of gene and intergenic length we used the more complete assembly for *P. striiformis* f. sp. *tritici*_104 E137A, previously excluded from orthologue analysis due to diploidy andexcluded the strain *P. striiformis* 93-210. We used custom R scripts to determine gene length and intergenic length distribution across datasets to determine alternative explanations for genome expansion. Untranslated regions (UTR) were not annotated for every gene, therefore, the analysis includes UTRs.

## Availability of supporting data

### Genome assembly and annotation data files

The primary genome assembly file, named APSI_primary.v1.fasta, has been deposited at the European Nucleotide Archive (ENA) at EMBL-EBI under the following ENA accession: ERZ1194194 (GCA_902702905). Raw data is deposited at ENA under project: PRJEB34084 (Study accession ERP116938). Locus tags are registered as APSI (for *<u>A</u>ustropuccinia <u>psi</u>dii*) and scaffolds identified as APSI_Pxxx (where x indicates scaffold number) for primary assembly. Annotation data files and the di-haploid assembly file (APSI_v1.fa) are available from DOI:10.5281/zenodo.3567172. The di-haploid assembly incorporates the 67 secondary assembly scaffolds (APSI_Hxxx for secondary). Assembly and annotation scripts as well as data files are available at https://github.com/peritob/Myrtle-rust-genome-assembly: including the draft mitochondrial sequence, APSI_primary_v1.xlsx and APSI_secondary_v1.xlsx (containing HMMER and BLAST against transcriptome results).

### Additional files

Additional Figure 1.png

The GC content profile as identified by OcculterCut. of the *M. brunnea, R. commune, A.psidii, P. graminis* f. sp. *tritici, P. triticina, P. coronata f. sp. avenae, P. sorghi, P. striifomis f. sp. tritici, C.quercuum* f. sp. *fusiforme* and *M. larici-populina* genome. AT-rich isochores or regions can be observed in *M.brunnea, R.commune* and *A. psidii*, but peaks of the first two fungal species are more separated than that of *A. psidii*. Also, compared to other fungi here, *A. psidii* has the overall higher AT-content. AT-rich isochores cannot be detected in all other seven Pucciniomycetes.

Additional Figure 2.html

APSI_primary and secondary_v1 genome and transcriptome mapping to the NCBI fungal database and visualised with Krona/2.7.

Additional Data 1.docx

Variant analysis of resequencing DNA from Australian isolates of *Austropuccinia psidii*

Additional Table 1.xlsx

(A) Annotation of rust genomes with 21 most abundant TE families using RepeatMasker. Initial annotation in the *A. psidii* genome was done using the REPET pipeline and might vary slightly from this annotation. The top 24 TE families were identified by deTEnovo from REPET. The top 24 repeats were identified previously and included in Repbase. (B) 5mC DNA MTases and 6mA DNA MTases annotations for *A. psidii*.

Additional Table 2.xlsx

Pucciniales comparative statistics, including Pfam analysis of tree rust specific gene orthologs, and Interproscan (v.5) results from the primary and secondary annotated assemblies.

Additional Table 3.xlsx

Hidden Markov model (HMMER) functional annotation of the predicted effectors within the primary assembly based on a reference proteome database (Potter et al. 2018). Listed are the 77 predicted effectors that had a functional annotation match.

## Abbreviations

BUSCO: Benchmarking Universal Single-Copy Orthologs; gDNA: genomic DNA; TE: transposable elements; PacBio: Pacific Biosciences; RNA-seq: RNA sequencing; SMRT: single-molecule real time (SMRT®); GB: Gigabyte; Gb: Gigabase

## Author’s contribution

**R.P.** conceived of, supported the study, provided the samples and made significant contributions to directing the research. **P.T.** and **B.S.** are shared first authors. **P.T.** did flow cytometry, assembled, annotated the genome, ran analyses and wrote much of the manuscript. **B. S.** made substantial efforts in directing the research, made Hi-C libraries, ran analyses including TEs and genome structure and helped write the manuscript. **C.D**. extracted and sequenced the gDNA and RNA. **C.H.D.** and **C.W.** ran Hi-C and other analyses. **A.J.** made Hi-C libraries. **K.S.** provided urediniospores, DNA of exotic isolates and ran inoculations for RNA. **J.S.** ran analyses on scaffolds and contributed to the manuscript. **Z.L.** ran TE analyses. **J.T.** provided mitochondrial sequence data and helped with RNA sequencing. **G.S.** with **D.C**. who also ran SNP analysis, secured financial support for the research, and helped to co-ordinate the study. **P.Z.** helped with short-read DNA sequencing. All authors read and approved the final manuscript.

## Acknowledgements

We wish to thank Mark Powrie who generously helped with computational support throughout the assembly and annotation process. The authors acknowledge the Sydney Informatics Hub and the University of Sydney’s High Performance Computing cluster, Artemis, for providing the computing resources that have contributed to the research results reported within this paper. Thanks also to Dr Geoff Pegg, Queensland Department of Agriculture Fisheries and Forestry, for sharing recent field observations on the impacts of myrtle rust. We also thank Mr. Andrew McIntosh for his technical support in the greenhouse work. We thank Simone Fouche for inspiring discussion about transposable elements in fungi.

