## Supplementary figures and images for "*Austropuccinia psidii*, causing myrtle rust, has a gigabase-sized genome shaped by transposable elements"

### Additional Figure 1

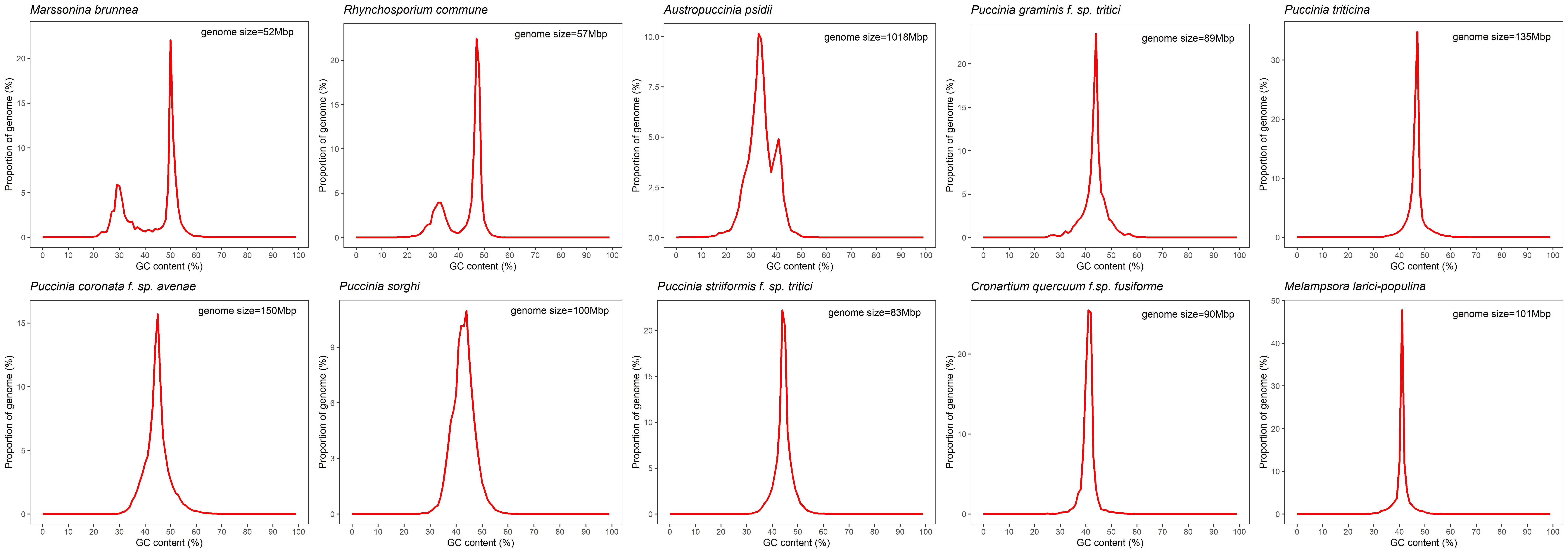
