## Additional Figure 2 for "*Austropuccinia psidii*, causing myrtle rust, has a gigabase-sized genome shaped by transposable elements": Additional Figure 1.HTML

Javascript must be enabled to view this page.

members
magnitude
magnitudeUnassigned
count
unassigned
taxon
rank

APSI\_primary
APSI\_secondary
TranscriptomeRepGenes
TranscriptomeAltGenes

666777682105014

node1.members.0.jsnode1.members.1.jsnode1.members.2.jsnode1.members.3.js
23217138895141

2759
434662949873
superkingdom

4751
node3.members.2.jsnode3.members.3.js
66
kingdom
434662949873

phylum
76
6029

76
suborder
6032

36734
76
family

6033
genus
76

species
34
node8.members.2.jsnode8.members.3.js
58839

node9.members.2.jsnode9.members.3.js
6035
32
species

species
1
node10.members.2.js
571949

node11.members.1.jsnode11.members.2.jsnode11.members.3.js
451864
434662819861
subkingdom
16681150

node12.members.2.jsnode12.members.3.js
5204
phylum
51304439
21

2
subphylum
51229323
452284
node13.members.2.js

5257
51206300
class

51206300
order
5267

1015
family
51206300
5268
node16.members.2.jsnode16.members.3.js

genus
51103147
63265

species
51103147
280036
node18.members.0.jsnode18.members.1.jsnode18.members.2.jsnode18.members.3.js

genus
93138
5269

5270
node20.members.2.jsnode20.members.3.js
93138
species

1538075
class
2123

2123
order
162474

2123
family
742845

55193
2123
genus

species
2123
node25.members.2.jsnode25.members.3.js
76775

5302
subphylum
73115

73115
class
155616

5234
73115
order

73115
family
1884633

116
genus
73115
5206
node30.members.2.jsnode30.members.3.js

1884637
species group
2350

species
2350
node32.members.2.jsnode32.members.3.js
37769

3959
species group
1897064

5207
node34.members.2.jsnode34.members.3.js
56
species
3959

178876
node35.members.2.jsnode35.members.3.js
2220
varietas

40410
node36.members.2.jsnode36.members.3.js
1233
varietas

384453098272
phylum
17151087
node37.members.1.jsnode37.members.2.jsnode37.members.3.js
4890

subphylum
273114511741
147537

4891
class
273114511741

order
273114511741
143125
node40.members.2.jsnode40.members.3.js
4892

2333
family
115784

460517
genus
2333

2333
species
460519
node43.members.2.jsnode43.members.3.js

node44.members.2.jsnode44.members.3.js
766764
family
611427520
43

genus
13642
4958

4959
species
13642

varietas
13642
node47.members.0.jsnode47.members.2.jsnode47.members.3.js
58641

511362443
genus
1516
node48.members.2.jsnode48.members.3.js
1535326

47187214
species
42374
node49.members.0.jsnode49.members.1.jsnode49.members.2.jsnode49.members.3.js

node50.members.1.jsnode50.members.2.jsnode50.members.3.js
5476
4117131
species

273371
node51.members.0.jsnode51.members.2.jsnode51.members.3.js
14382
species

766733
genus
2532

species
2532
4924
node53.members.2.jsnode53.members.3.js

family
3275125
34353

genus
3275125
4951

species
3275125
node56.members.0.jsnode56.members.1.jsnode56.members.2.jsnode56.members.3.js
4952

1156497
family
735472

461281
918
genus

node59.members.2.jsnode59.members.3.js
1005962
species
918

4919
genus
734554

species
734554
4909
node61.members.0.jsnode61.members.1.jsnode61.members.2.jsnode61.members.3.js

410830
family
4778

genus
4778
410829

4778
species
node64.members.2.jsnode64.members.3.js
796027

4893
node65.members.0.jsnode65.members.2.jsnode65.members.3.js
14753
family
1115682788

612145114
genus
1
node66.members.2.js
113604

species
61110678
1071379
node67.members.0.jsnode67.members.1.jsnode67.members.2.jsnode67.members.3.js

node68.members.1.jsnode68.members.2.jsnode68.members.3.js
113608
species
13836

21
genus
5674
33170
node69.members.2.jsnode69.members.3.js

45285
node70.members.2.jsnode70.members.3.js
species
2536

33169
node71.members.2.jsnode71.members.3.js
1314
species

1623
species
45286
node72.members.2.jsnode72.members.3.js

300275
1518
genus

381046
node74.members.2.jsnode74.members.3.js
1518
species

11121140
genus
1
node75.members.2.js
278028

node76.members.0.jsnode76.members.1.jsnode76.members.2.jsnode76.members.3.js
27289
118695
species

species
3445
node77.members.2.jsnode77.members.3.js
27288

genus
1221
4948

1221
species
node79.members.2.jsnode79.members.3.js
4950

318889
genus
4910

species
316171
node81.members.0.jsnode81.members.1.jsnode81.members.2.jsnode81.members.3.js
4911

species
2718
28985
node82.members.2.jsnode82.members.3.js

genus
2645
374468

2645
species
5478
node84.members.2.jsnode84.members.3.js

genus
12527
4953

12527
species
node86.members.1.jsnode86.members.2.jsnode86.members.3.js
4956

4930
node87.members.2.js
2
7599
genus

species
3334
4932
node88.members.2.jsnode88.members.3.js

species
4065
1080349
node89.members.2.jsnode89.members.3.js

72108
genus
71245

node91.members.2.jsnode91.members.3.js
588726
2236
species

5072
species
432096
node92.members.2.jsnode92.members.3.js

451866
14347
subphylum

14347
class
147554

order
14347
34346

4894
14347
family

14347
genus
4895

14347
species
4896
node98.members.0.jsnode98.members.2.jsnode98.members.3.js

291482
101231005397
subphylum
147538
node99.members.2.jsnode99.members.3.js

147541
class
1176135

1176135
subclass
451867

134362
1176135
order

node103.members.3.js
93133
1176135
family
1

1047167
genus
115598

node105.members.0.jsnode105.members.1.jsnode105.members.2.jsnode105.members.3.js
1047171
species
115598

genus
2136
29002

species
2136
122368
node107.members.2.jsnode107.members.3.js

81023544165
class
1284499
node108.members.1.jsnode108.members.2.jsnode108.members.3.js
147550

5593
1311461874
subclass
222544
node109.members.2.jsnode109.members.3.js

13366557
order
639021

81093
13366557
family

genus
119201
148303

node113.members.2.jsnode113.members.3.js
318829
species
119201

node114.members.2.jsnode114.members.3.js
48558
13247356
genus
29

1578925
node115.members.0.jsnode115.members.1.jsnode115.members.2.jsnode115.members.3.js
13245347
species

7251224
order
2941
node116.members.2.jsnode116.members.3.js
5139

1841
497850
family
35718
node117.members.2.jsnode117.members.3.js

35719
124218
genus

species
124218
node119.members.2.jsnode119.members.3.js
35720

1920207
genus
355591

78579
node121.members.2.jsnode121.members.3.js
species
355591

family
199333
5148

genus
199333
5140

node124.members.2.jsnode124.members.3.js
5141
species
199333

769241792
subclass
4066
node125.members.2.jsnode125.members.3.js
222543

1028384
order
5291028

681950
5291028
family

genus
5291028
5455

node129.members.2.jsnode129.members.3.js
80884
species
5291028

5125
node130.members.2.jsnode130.members.3.js
615
order
76355698

34397
family
4368

243023
genus
4368

species
4368
280754
node133.members.2.jsnode133.members.3.js

family
76306615
110618

1555
genus
76306615
5506
node135.members.2.jsnode135.members.3.js

171631
1147100
species group

1147100
species
5507
node137.members.0.jsnode137.members.1.jsnode137.members.2.jsnode137.members.3.js

569360
node138.members.2.jsnode138.members.3.js
1548
species group
65131248

node139.members.2.jsnode139.members.3.js
5518
3565
species

14080
species
node140.members.0.jsnode140.members.2.jsnode140.members.3.js
56646

species
554155
node141.members.0.jsnode141.members.1.jsnode141.members.2.jsnode141.members.3.js
101028

411
species group
113212
171627
node142.members.2.jsnode142.members.3.js

node143.members.2.jsnode143.members.3.js
117187
species
62114

species
4787
5127
node144.members.2.jsnode144.members.3.js

147548
1266405
class

5178
order
1266405

28983
family
1266405

genus
1266405
33196

40559
node149.members.1.jsnode149.members.2.jsnode149.members.3.js
1266405
species

class
1113210
147545

subclass
1113210
451871

5042
1113210
order

1131492
1113210
family

22
genus
1113210
5052
node154.members.2.jsnode154.members.3.js

59124
species
5062
node155.members.2.jsnode155.members.3.js

746128
node156.members.0.jsnode156.members.2.jsnode156.members.3.js
15284
species
