## Additional Data 1 for "*Austropuccinia psidii*, causing myrtle rust, has a gigabase-sized genome shaped by transposable elements"

### Methods

#### Variant analysis of Australian isolates

Whole genome DNA resequencing on the Hiseq Illumina sequencing platform (250 bp paired-end reads) was obtained from six other Australian isolates, one isolate from Hawaii believed to be from the pandemic strain, and a Brazilian isolate stored at the University of Sydney, Plant Breeding Institute (PBI). A total of 209 GB of (250 bp) paired-end sequence data was first trimmed and then Bowtie2 (Langmead et al. 2009) was used for mapping reads to the primary genome assembly using the --end-to-end option for each sample independently. The sam file of each sample was then converted into bam files using Samtools (Li et al. 2009), sorted and indexed against the reference genome. Samtools mpileup and bcftools were used to generate compressed variant calling files (vcf.gz). Variants were filtered using vcftools to a minimum depth of 6X and insertion-deletions were removed. The vcf file was sorted and format converted with Tassel v5 (Bradbury et al. 2007) and the phylip alignment file was run with iqtree v1.6.7 (Nguyen et al. 2015) with parameters -bb 1000 -nt AUTO and the Brazilian isolate as outgroup. Phylogenetic evolutionary relationships based on 44,375 SNPs within pathogen isolates collected in Australia (Au), Hawaii (Hw) and outgroup from Brazil (Bz, from *Eucalyptus grandis*) were visualized with Dendroscope (v.3) (Huson & Scornavacca 2012).

### Results

#### Pandemic isolate variation identified in Australia

The DNA from these isolates was previously tested using simple sequence repeat markers and all Australian isolates were determined to be genetically similar (Sandhu et al. 2015). When comparing pathogen isolates, we found 44,375 SNPs in total. On average one SNP per 23 Kbp. The maximum likelihood phylogenetic tree, in Newick format, was visualized in Dendroscope v3 (Huson & Scornavacca 2012) (Figure A) and indicates variations present in the pandemic strain of the pathogen. These isolates were collected from diverse hosts and locations in Australia dating back to 2011 (Table A). Isolate Au 7, collected from the host plant *Melaleuca quinquenervia* in 2012, Queenscliff, NSW, could not be separated from the Brazilian isolate with strong bootstrap values. Nevertheless, the clustering away from other Australian isolates is concerning and perhaps indicates that a variant is already present in Australia.

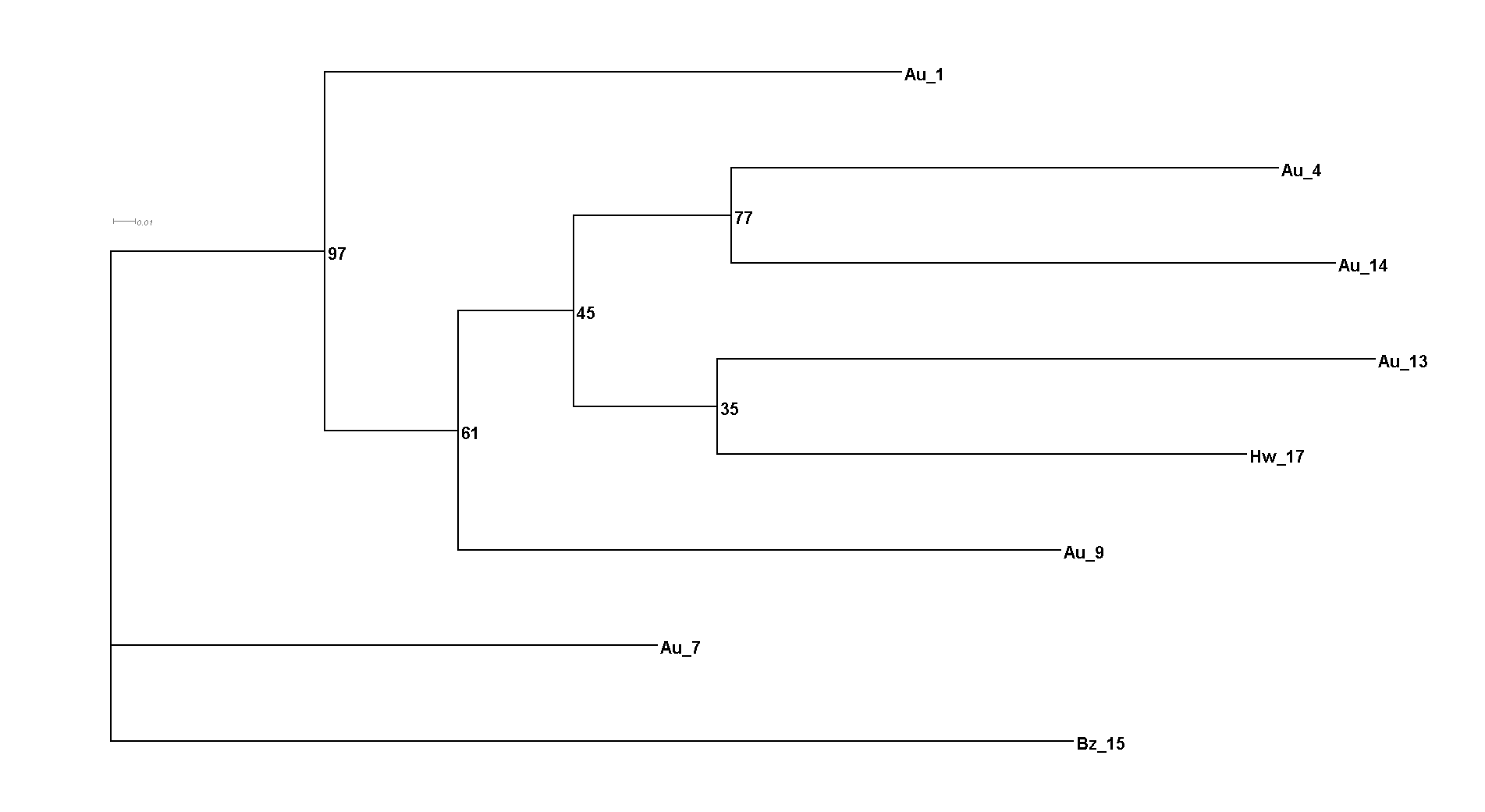

Fig. A. Phylogenetic evolutionary relationship based on 44,375 single nucleotide polymorphisms within pathogen isolates collected in Australia (Au), Hawaii (Hw) and outgroup from Brazil (Bz from *Eucalyptus grandis*) inferred using the Maximum likelihood method with Iqtree v1.6.7 and visualized with Dendrosope v3. The tree is UNROOTED although outgroup taxon Bz is drawn at root. Numbers are standard bootstrap support (%). Au 7 and Bz could not be separated with strong bootstrap values. All other isolates cluster with Hw but bootstrap values between sub-branches are weak.

Table A. Details of local and exotic isolates of *Austropuccinia psidii* maintained at the University of Sydney, Plant Breeding Institute (PBI), and used for preliminary variant analysis. All Au isolates were increased from single pustule isolate on *Syzygium jambos* (Sandhu et al. 2015) .

| **Isolate ID** | **Acc. No.** | **Original host** | **Location** | **Year of Collection** |
| --- | --- | --- | --- | --- |
| Au_1 | 115001 | *Syzygium jambos* | Lismore, NSW | 2011 |
| Au_4 | 125004 | *Eucalyptus pilularis* | Newry, NSW | 2012 |
| Au_7 | 125009 | *Melaleuca quinquenervia* | Queesncliff, NSW | 2012 |
| Au_9 | 125014 | *Chamelaucium uncinatum* | Toowoomba, QLD | 2012 |
| Au_13 | 135001 | *Rhodamnia maideniana* | Mooball, NSW | 2013 |
| Au_14 | 135002 | *Rhodamnia rubescens* | Nightcap N. P., NSW | 2013 |
| Hw_17 | 135007 | *Syzygium jambos* | Hawaii, USA | 2013 |
| Bz_15 | 135005 | *Eucalyptus grandis* | Vicosa, Brazil | 2013 |
