## Additional Table 3 for "*Austropuccinia psidii*, causing myrtle rust, has a gigabase-sized genome shaped by transposable elements"

#### Hidden Markov model (HMMER) functional annotation of the predicted effectors within the primary assembly based on a reference proteome database (Potter et al. 2018). Listed are the 77 predicted effectors that had a functional annotation match.

| **Query Name** | **Hits** | **Top Hit** | **Details** | **E-value** |
| --- | --- | --- | --- | --- |
| **APSI_P001.5595.t2** | **1** | **A0A371AUW5_9FIRM** | **Peptidase S1** | **0.0021** |
| **APSI_P001.6256.t1** | **3** | **G3AVB1_SPAPN** | **Candida_ALS_N domain-containing protein** | **3.30E-07** |
| **APSI_P001.6257.t1** | **3** | **G3AVB1_SPAPN** | **Candida_ALS_N domain-containing protein** | **3.30E-07** |
| **APSI_P001.6259.t1** | **3** | **G3AVB1_SPAPN** | **Candida_ALS_N domain-containing protein** | **3.30E-07** |
| **APSI_P001.6338.t1** | **1158** | **E3KKC7_PUCGT** | **Succinate dehydrogenase [ubiquinone] cytochrome b small subunit** | **2.10E-66** |
| **APSI_P001.6438.t1** | **5** | **A0A1S3F6G5_DIPOR** | **zinc finger protein 260-like** | **0.00015** |
| **APSI_P001.6823.t1** | **1375** | **A0A0C4DFA7_PUCT1** | **60S ribosomal protein L29** | **5.10E-26** |
| **APSI_P001.6864.t1** | **5156** | **A0A2N5UW88_9BASI** | **Small nuclear ribonucleoprotein G** | **1.30E-41** |
| **APSI_P001.7258.t1** | **3667** | **A0A0L0W3I3_9BASI** | **Tyrosinase_Cu-bd domain-containing protein** | **2.70E-143** |
| **APSI_P002.14561.t1** | **506** | **A0A2S4UMX1_9BASI** | **Lipase_3 domain-containing protein** | **6.40E-89** |
| **APSI_P002.14561.t2** | **582** | **A0A0L0V6Q5_9BASI** | **Lipase_3 domain-containing protein** | **4.40E-103** |
| **APSI_P002.14778.t1** | **3** | **D5STC2_PLAL2** | **YD repeat protein** | **4.40E-06** |
| **APSI_P002.15786.t1** | **5** | **A0A0C3FWT6_9AGAM** | **WD_REPEATS_REGION domain-containing protein (Fragment)** | **0.00013** |
| **APSI_P002.15789.t1** | **2** | **S8FCT5_9BACT** | **PF03382 family protein** | **0.00055** |
| **APSI_P003.1876.t1** | **2** | **A9VDE7_MONBE** | **Predicted protein** | **4.30E-06** |
| **APSI_P003.1934.t1** | **67** | **E3JSA4_PUCGT** | **Lipase_3 domain-containing protein** | **1.30E-121** |
| **APSI_P003.2310.t1** | **2** | **A0A4C1T5X5_9NEOP** | **Titin (Fragment)** | **2.30E-18** |
| **APSI_P003.2387.t1** | **3** | **A0A287AE45_PIG** | **PDZ domain-containing protein** | **7.30E-18** |
| **APSI_P003.2523.t1** | **520** | **A0A2N5V3I2_9BASI** | **Chorismate mutase domain-containing protein** | **7.90E-65** |
| **APSI_P004.2961.t1** | **262** | **F4R7G1_MELLP** | **Beta_helix domain-containing protein** | **7.40E-87** |
| **APSI_P004.3404.t1** | **1** | **A7T525_NEMVE** | **Predicted protein (Fragment)** | **0.00025** |
| **APSI_P004.3462.t1** | **1480** | **F4RU02_MELLP** | **Family 5 carbohydrate esterase** | **9.60E-49** |
| **APSI_P004.3557.t1** | **2457** | **A0A180G9U2_PUCT1** | **Mannan endo-1,6-alpha-mannosidase** | **3.50E-129** |
| **APSI_P005.10407.t1** | **2** | **A0A210PML9_MIZYE** | **Enterin neuropeptide** | **3.60E-06** |
| **APSI_P005.10417.t1** | **2** | **A0A210PML9_MIZYE** | **Enterin neuropeptide** | **7.20E-06** |
| **APSI_P005.10459.t1** | **193** | **F4RV95_MELLP** | **Secreted protein** | **3.90E-56** |
| **APSI_P005.11212.t1** | **1934** | **A0A0C4F208_PUCT1** | **S10_plectin domain-containing protein** | **6.40E-86** |
| **APSI_P006.10022.t1** | **1** | **W4ZC84_STRPU** | **ANK_REP_REGION domain-containing protein** | **0.00049** |
| **APSI_P006.9367.t1** | **1044** | **A0A0B7FB57_THACB** | **SPBc2 prophage-derived uncharacterized transglycosylase YomI** | **2.40E-116** |
| **APSI_P006.9804.t1** | **807** | **F4R776_MELLP** | **Secreted protein** | **2.30E-44** |
| **APSI_P007.13556.t1** | **6** | **A0A166LR54_9PEZI** | **Kinesin light chain 3 (Fragment)** | **4.80E-06** |
| **APSI_P007.14266.t1** | **3704** | **V2XK15_MONRO** | **Non-catalytic module family expn protein** | **3.10E-70** |
| **APSI_P007.14386.t1** | **163** | **F4RBM4_MELLP** | **CFEM domain-containing protein** | **9.40E-12** |
| **APSI_P007.14430.t1** | **2** | **A0A1S3WXY1_TOBAC** | **extensin-2-like** | **0.00014** |
| **APSI_P008.16742.t1** | **4862** | **A0A180G6Z8_PUCT1** | **SurE domain-containing protein** | **1.20E-105** |
| **APSI_P008.16775.t1** | **2** | **A0A1W2H7R2_9BACT** | **Conserved repeat domain-containing protein** | **0.003** |
| **APSI_P008.17031.t1** | **125** | **A0A2S4V7D1_9BASI** | **DH domain-containing protein** | **4.90E-39** |
| **APSI_P008.17121.t1** | **156** | **F4RJ46_MELLP** | **Secreted protein** | **4.90E-42** |
| **APSI_P009.17593.t1** | **5809** | **A0A1B0F9P6_GLOMM** | **CG16995** | **1.30E-87** |
| **APSI_P010.11572.t1** | **1** | **B0JGP8_MICAN** | **WD-repeat protein** | **0.0054** |
| **APSI_P010.12024.t1** | **2** | **A7T0Q8_NEMVE** | **Predicted protein (Fragment)** | **0.00034** |
| **APSI_P010.12049.t1** | **1** | **A0A091DWW5_FUKDA** | **T-cell surface glycoprotein CD1b** | **0.0058** |
| **APSI_P011.414.t1** | **568** | **F4R776_MELLP** | **Secreted protein** | **5.80E-38** |
| **APSI_P011.451.t1** | **733** | **F4R776_MELLP** | **Secreted protein** | **5.40E-41** |
| **APSI_P011.466.t1** | **927** | **F4R776_MELLP** | **Secreted protein** | **2.60E-51** |
| **APSI_P011.649.t1** | **25116** | **F4RTN7_MELLP** | **GHMP_kinases_N domain-containing protein** | **9.20E-85** |
| **APSI_P011.659.t1** | **4054** | **V2XK15_MONRO** | **Non-catalytic module family expn protein** | **1.20E-85** |
| **APSI_P012.8825.t1** | **11** | **A0A067MSE4_9AGAM** | **WD_REPEATS_REGION domain-containing protein (Fragment)** | **2.90E-07** |
| **APSI_P012.9030.t1** | **1** | **D1P3X8_9GAMM** | **M protein repeat protein (Fragment)** | **6.30E-09** |
| **APSI_P012.9205.t1** | **136** | **A0A0L0V0D2_9BASI** | **SET domain-containing protein** | **1.10E-73** |
| **APSI_P012.9228.t1** | **3** | **A7S3T5_NEMVE** | **Predicted protein (Fragment)** | **0.00036** |
| **APSI_P013.3906.t1** | **76** | **A0A2S4V583_9BASI** | **Integrase catalytic domain-containing protein** | **8.10E-20** |
| **APSI_P013.3907.t1** | **3** | **A0A3B0CJ79_9BACL** | **DUF5018 domain-containing protein** | **4.00E-06** |
| **APSI_P013.4035.t1** | **4** | **A0A4V2T0V8_9PAST** | **Filamentous hemagglutinin (Fragment)** | **0.00046** |
| **APSI_P013.4080.t1** | **411** | **E3JXI7_PUCGT** | **Long chronological lifespan protein 2** | **9.10E-22** |
| **APSI_P013.4092.t3** | **2618** | **A0A2N5U9K5_9BASI** | **GPI-anchor transamidase** | **3.30E-84** |
| **APSI_P013.4169.t1** | **2** | **A0A498MYR2_LABRO** | **Protocadherin alpha-C2-like protein** | **0.00092** |
| **APSI_P013.4384.t1** | **2** | **A7T7U2_NEMVE** | **Predicted protein (Fragment)** | **0.00022** |
| **APSI_P015.12851.t1** | **2** | **E9AD26_LEIMA** | **Putative calpain-like cysteine peptidase** | **2.80E-22** |
| **APSI_P016.16039.t1** | **1** | **A9UUZ3_MONBE** | **Predicted protein** | **0.0031** |
| **APSI_P016.16116.t1** | **1** | **A7S9R6_NEMVE** | **Predicted protein** | **2.20E-10** |
| **APSI_P016.16116.t2** | **1** | **A7S9R6_NEMVE** | **Predicted protein** | **1.80E-10** |
| **APSI_P016.16126.t1** | **11111** | **A0A1Y2E9I9_9PEZI** | **Alpha/Beta hydrolase protein** | **1.10E-77** |
| **APSI_P016.16246.t1** | **1** | **A0A1S3QJL1_SALSA** | **endo-1,4-beta-xylanase B-like** | **4.50E-12** |
| **APSI_P016.16247.t1** | **1** | **A0A1S3QJL1_SALSA** | **endo-1,4-beta-xylanase B-like** | **5.20E-12** |
| **APSI_P017.12604.t1** | **3** | **A8DWD8_NEMVE** | **Predicted protein (Fragment)** | **6.30E-05** |
| **APSI_P018.7870.t1** | **1895** | **A0A0D2X116_CAPO3** | **SH3 domain-containing protein** | **2.30E-116** |
| **APSI_P019.8195.t1** | **1589** | **A0A180GGI2_PUCT1** | **PRELI/MSF1 domain-containing protein** | **5.20E-62** |
| **APSI_P019.8218.t1** | **8019** | **A0A0L6UB46_9BASI** | **Phenylalanine--tRNA ligase** | **5.40E-61** |
| **APSI_P019.8306.t1** | **1** | **A0A1V9Y7C9_9STRA** | **Major Facilitator Superfamily (MFS)** | **0.00086** |
| **APSI_P019.8320.t1** | **1** | **A7RGU2_NEMVE** | **Predicted protein** | **0.00049** |
| **APSI_P021.13316.t1** | **2** | **M7BTP8_CHEMY** | **Nucleolar RNA helicase 2 (Fragment)** | **0.0013** |
| **APSI_P022.4675.t1** | **3** | **A7RJS0_NEMVE** | **Predicted protein** | **0.00015** |
| **APSI_P022.4680.t1** | **5** | **F4RG77_MELLP** | **Secreted protein** | **1.90E-06** |
| **APSI_P022.4793.t1** | **1** | **A0A1D3TYV3_9FIRM** | **LPXTG-motif cell wall anchor domain-containing protein** | **0.0053** |
| **APSI_P023.8485.t1** | **3** | **A0A0R4IE08_DANRE** | **Si:dkey-28k24.2** | **0.00067** |
| **APSI_P024.16551.t1** | **1** | **A0A1P8F4T7_9CHLR** | **Cna protein B-type domain-containing protein** | **5.50E-07** |
